# Prediction of *α*_*IIb*_*β*_3_ integrin structures along its minimum free energy activation pathway

**DOI:** 10.1101/2025.05.16.654557

**Authors:** Siva Dasetty, Robert E. Coffman, Tamara C. Bidone, Andrew L. Ferguson

## Abstract

The adhesion protein integrin is a transmembrane heterodimer that plays a pivotal role in cellular processes such as cell signaling and cell migration. To execute its function, integrin undergoes extensive conformational changes from a bent-closed to an extended-open state. Resolving the structures across these changes remains a challenge with both experimental and computational methods, but is crucial for understanding the activation mechanism of integrin. We address this challenge for the platelet integrin *α*_*IIb*_ *β*_3_ by employing finite temperature string method with structures of the images along the initial guess path generated by a multiscale data-driven framework. The full-length all-atom structures along the resulting minimum free energy path between the inactive bent-closed and active extended-open states of *α*_*IIb*_ *β*_3_ integrin are consistent with a variety of experimentally resolved structures. Changes in these predicted structures along the path show that the extension and separation of the *α* and *β* subunits from the bent-closed to the extended-open state require correlated movements between the subdomain pairs in *α*_*IIb*_ *β*_3_. These results provide new insights into integrin activation mechanism and the predicted structures have potential applications in guiding the design of integrin targeting therapeutics.

**SIGNIFICANCE:** Integrins are receptor proteins that mediate a number of critical cellular processes, but it remains challenging to resolve the molecular details of integrin activation by both experiments and simulations. We address this challenge by employing a computational method to study rare events and resolve the all-atom transient structures of a platelet integrin, *α*_*IIb*_ *β*_3_. The resulting structures are in good agreement with experimentally resolved partial structures and additionally reveal correlated movements between the *α*_*IIb*_ *β*_3_ integrin subdomain pairs during its transition from the inactive to the active conformational state. The structures predicted in this work have applications as therapeutic targets to address diseases linked to *α*_*IIb*_ *β*_3_ integrin dysfunction, such as bleeding and thrombotic disorders.

## INTRODUCTION

Integrin is a large transmembrane heterodimer protein consisting of non-covalently associated *α* and *β* subunits.(1) It plays a critical role in several cellular functions such as cell signaling, adhesion, and differentiation by responding to signals both within and outside the cell.(2–4) There are 18 *α* and 8 *β* subunits that together form the 24 known variants of integrins in humans.(4, 5) The *α* subunit is comprised of ∼ 940 to 1120 residues, depending on the variant, with the domains from N- to C-terminal referred to as *β*_*propellar*_ , thigh, calfs (calf-1, calf-2), and a transmembrane helix (*α*_*helix*_).(6–8) In contrast, *β* subunit has ∼ 700 residues and is formed by the domains, from N-terminus to C-terminus, *β*_*A*_, hybrid, plexin-semaphorin-integrin (PSI), epidermal growth factor (EGF-I) modules E1-E4, *β*_*tail*_ (*β*_*T*_ ), and a transmembrane helix (*β*_*helix*_ .(6–8) The *β*_*propellar*_ and *β*_*A*_ domains, respectively, form the ectodomain head parts of *α* and *β* subunits, which contain the extra-cellular ligand binding site at their contact interface. The thigh and two calf domains are known as the leg region of the *α* subunit. In the *β* subunit, the leg region is formed by the hybrid domain, PSI, EGF-I module, and *β*_*T*_ domain (*β*_*TD*_). Both the *α* and *β* subunits contain a flexible region between the head and leg regions. In the *α* subunit, this region is called the genu and it exists between the thigh and calf-1 subdomains. The flexible linkers are not well resolved in the *β* subunit but are approximately located within the EGF-I domains, and between the PSI and hybrid domain(4). Fig. 1 shows an illustration of all of these domains in the platelet integrin, *α*_*IIb*_ *β*_3_.

**Figure 1:**
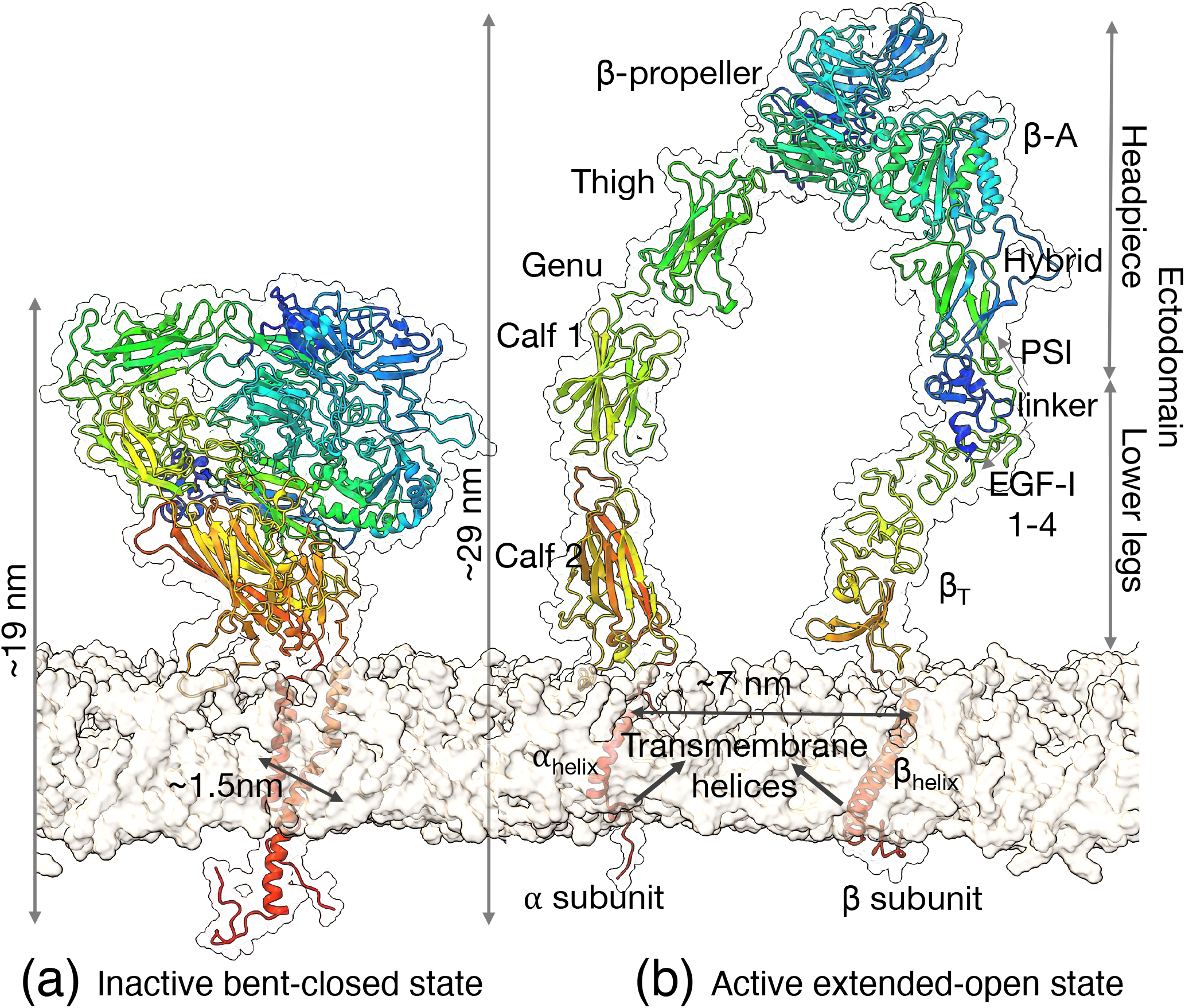
Illustration of the key all-atom structures of *α*_*IIb*_ *β*_3_ integrin along its activation pathway. (a) Low-affinity structure corresponding to the bent-closed state. (b) High-affinity structure of the extended-open state. The reversible large-scale conformational change between the low-affinity and high-affinity structures can be triggered bi-directionally either through inside-out or outside-in activation. The transition involves separation of the transmembrane helices and extension of the closely associated ectodomains of the *α* and *β* subunits at the genu or linker regions. Both the protein structures in cartoon representation are rendered with lipid bilayer using UCSF ChimeraX 1.6.1(13, 14).

In the inactive state, integrin adopts a low-affinity conformation known as the bent-closed state, where the ectodomains of both the *α* and *β* subunits are closely associated and bent toward the cell membrane at the flexible genu and linker regions (Fig. 1a).(9) When activated by intracellular proteins (“inside-out” activation) or extracellular ligands (“outside-in” activation), integrin transitions reversibly on the millisecond(10)-second(11) timescale into a high-affinity active conformation called the extended-open state (Fig. 1b). This involves extension of the ectodomains of both *α* and *β* subunits into an upright conformation and separation of the transmembrane helical regions by ∼ 7-8 nm.(4, 9) Resolving the structural pathway of this conformational change is critical for understanding the function of integrin in cellular interactions and in guiding the development of therapeutics for diseases associated with its malfunction or its over-expression in cancer cells.(7, 12)

Several potential structural pathways for integrin activation by both inside-out and outside-in signals have been postulated.(6, 9, 10, 12, 15, 16) Notable examples are the switchblade(6), deadbolt(16), and concerted activation(10) mechanism models. According to the switchblade model, the extended-open state is realized from the bent-closed state first by a separation of the legs and transmembrane helices followed by unbending of the headpieces in *α* and *β* subunit, which in a motion akin to the opening of a pocket knife, exposes the ligand binding site to the extracellular matrix.(6, 12) As the name suggests, the deadbolt model refers to a mechanism involving the removal of a deadbolt from its lock. Specifically, first inside-out signalling results in a progressive loss of interactions between the *β*_*TD*_ and the *β*_*A*_ domains that act as the deadbolt and lock regions, respectively, in the bent-closed state.(16, 17) Following the release of the deadbolt, the legs are separated to fully transition into the extended-open state.(12, 17) In contrast to the independent movements of legs and headpieces in switchblade and deadbolt models, the concerted mechanism model posits that the leg separation and headpiece opening proceed in a concerted manner.(10)

Resolving the all-atom (AA) structures of integrin along the activation pathway using experimental methods is challenging due to the transient nature of the millisecond(10) to second(11) activation process. Molecular modeling can help shed light on the all-atom details of integrin activation, but unbiased molecular dynamics (MD) simulations are rendered extremely computationally expensive for large molecular systems such as integrin that contain ∼ 1710-1840 residues or more than 1M atoms with lipid bilayer and an explicit water model.(7, 18) Moreover, it may be unfeasible to cross the thermodynamic barriers separating the bent-closed and extended-open conformational states on timescales accessible to unbiased atomistic MD simulations. To address this, a non-equilibrium technique called steered MD simulations is often employed, wherein artificial forces are added to drive the structure from one state to another state.(12, 19, 19–22) While insights from these works are invaluable in finding critical domains or interactions governing the transition, the application of artificial bias forces could distort the structures to unfavorable transition paths under finite sampling times. Collective variable (CV) based enhanced sampling methods(23, 24) may also be applied to drive the activation transition, but to be effective require *a priori* knowledge of the transition paths for driving the sampling from one state to another state. While discriminatory CVs such as linear discriminant analysis (LDA)(25, 26) can tackle this problem, it can be challenging to apply them to large multi-molecular systems because of the computational cost associated with the system size. Similar computational cost associated issues exist for tempering based enhanced sampling methods such as replica exchange MD(27) and multi-canonical simulations(28). Coarse-grained (CG) modeling techniques can address these problems but their success is often limited due to challenges in parameterizing CG potentials that accurately capture the emerging dynamics along the activation pathway and additionally, coarse-graining results in an inherent loss in the atomistic resolution of the system.(11, 29, 30)

To address the above computational challenges, we have previously developed a multi-scale data-driven framework that integrates MD simulations, non-linear manifold learning, and deep generative modeling.(31) Conceptually, our approach learns a low-dimensional projection of the ensemble of experimentally known metastable states and then generates plausible activation pathways by interpolating in the learned low-dimensional latent space. In our previous work(31), we applied this generic framework to predict the transient structures of the platelet integrin, *α*_*IIb*_ *β*_3_, along the hypothesized switchblade, deadbolt, and concerted activation mechanism pathways. However, as extrapolative predictions, it remains unclear whether these predicted structures lie along the thermodynamically most favorable pathway. In this work, our goal is to address this question by finding the minimum free energy activation pathway and render the full-length all-atom structures along it. To achieve this, we apply the finite temperature string method (FTS)(32) with structures along the initial pathway generated using the approach developed in our prior work(31). We seed the FTS method with an initial path by following a hypothetical deadbolt mechanism. We found that the full-length AA structures along the relaxed pathway from the FTS method have a root mean square deviation (RMSD) of less than 1 nm with many of the partial transition structures resolved using X-ray crystallography, nuclear magnetic resonance (NMR) spectroscopy, and cryo-electron microscopy (cryo-EM) methods. The good experimental agreement lends confidence to the proposed minimum free energy activation pathway, which can shed new light on the molecular details of activation and identifies intermediate conformations that may be useful in guiding therapeutic development.

The structure of the remainder of the paper is as follows. First, we describe our computational workflow along with details on each module by starting with the construction of the initial guess pathway for the FTS method. We then present the relaxation trends of the pathway along with their convergence analysis. Subsequently, we validate the relaxed pathway by comparing the structures along the relaxed string or the minimum free energy pathway with available experimental structures. We follow this with a discussion on the potential transition mechanism from bent-closed state to extended-open state by focusing on the structural changes between the different domains of *α*_*IIb*_ *β*_3_ integrin along the relaxed pathway. Finally, we provide our perspective in quantifying the free energy landscapes along the proposed minimum free energy pathway using enhanced sampling methods and discuss the potential applications of the predicted structures in designing therapeutics to treat integrin dysfunction or malfunction.

## METHODS

Our computational approach consists of four components — (a) definition of an initial guess for a minimum free energy deadbolt activation pathway, (b) relaxation of this path using the FTS method, (c) exploration of the configurational space along the path using AA MD simulations, and (d) convergence assessment of the paths. We iterate through steps (b), (c), and (d) until the path is fully relaxed. Fig. 2 illustrates the overall workflow. In the following sections, we provide the methodological details of each component of the approach. We end this section by detailing the validation of the relaxed path by comparing against experimental structures of *α*_*IIb*_ *β*_3_ integrin.

**Figure 2:**
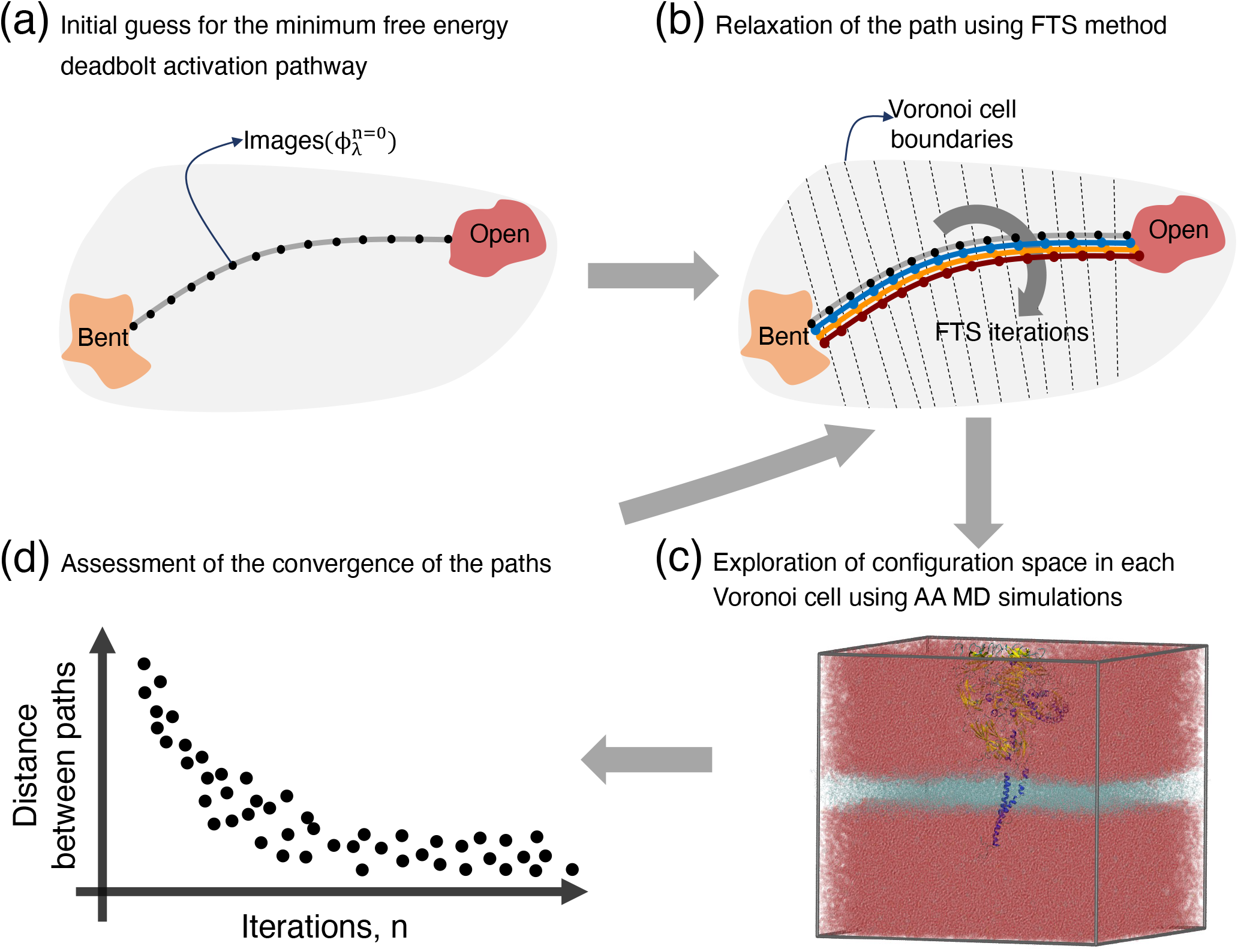
Graphical overview of the computational approach to predict the *α*_*IIb*_ *β*_3_ integrin activation pathway. (a) First, we guess an initial pathway (gray solid line) between the bent-closed state (orange) and the extended-open (red) state by tracing the deadbolt mechanism in a 2D physical CV space spanned by 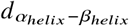 and 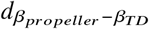, which represent the center of mass (COM) distances between *α*_*helix*_ and *β*_*helix*_, and *β*_*propellar*_ and *β*_*TD*_, respectively. We define the initial guess pathway as a string parameterized by *N*+1=19 equidistant points or images (black markers, 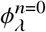), which we obtain by linear interpolation along an initial guess for the (unrelaxed) path generated using our previously reported approach(31) (SI Fig. S1). (b) Second, we relax the initial guess pathway to the minimum free energy pathway by iteratively employing the FTS method(32). In brief, we first determine the Voronoi boundaries (dashed lines) by using the images of the string in the current iteration as the nodes. Then, we estimate the images of the string in the next iteration (e.g., blue markers) by following the FTS method procedure. (c) We determine the configurational position of the system in each Voronoi cell in each iteration for updating the string according to FTS method by conducting 200 ps of AA MD simulations. Representative snapshot of the AA MD simulation system rendered using VMD 1.9.4(33). (d) We terminate the FTS method procedure by periodically (every 10 iterations) checking the convergence of the strings through the iterative cycle, and consider the final relaxed string as the minimum free energy pathway.

### Definition of the initial guess for the minimum free energy deadbolt activation pathway

We first define an initial guess path for the minimum free energy activation pathway (Fig. 2a) between the bent-closed and extended-open states in a 2D physical CV space spanned by 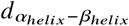 and 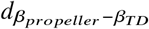. The 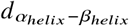 CV represents the COM distance between the transmembrane helices *α*_*helix*_ and *β*_*helix*_ in *α* and *β* subunits, respectively. 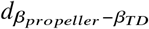 refers to the COM distance between the deadbolt (*β*_*propellar*_) and the lock domain (*β*_*TD*_) located in the *α* and the *β* subunits, respectively. Together 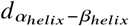 and 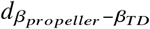 capture the key conformational differences in the ectodomain and transmembrane helices of the bent-closed and extended-open states of integrin. Furthermore, these two CVs can describe the deadbolt, switchblade, and concerted transition mechanisms of *α*_*IIb*_ *β*_3_ integrin (SI Fig. S1).(31) Specifically, the switchblade model can be imagined in a 2D space spanned by 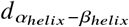 along x-axis and 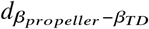along y-axis as first a horizontal transition in the x-axis followed by vertical transition in the y-axis. Conversely, the deadbolt model can be hypothesized as first a vertical transition in y-axis and then a horizontal transition in x-axis. A simultaneous transition along both x- and y-axis would correspond to the concerted mechanism model.

In this work, we focus on resolving the minimum free energy deadbolt activation pathway. We selected this path because two full-length extended intermediate structures of *α*_*IIb*_ *β*_3_ integrin, named Int1 and Int2, resolved using cryo-EM potentially lie along the deadbolt activation pathway, providing partial insights into its mechanism.(9, 18, 31) However, we note that we could perform analogous analyses to other initial paths, such as that of the switchblade or concerted mechanism models, within the framework developed in this work. In the Int1 structure, the ectodomain extends relative to the bent-closed structure at the flexible genu or linker regions. In addition to these changes, the hybrid domain in the *β* subunit swings out, and the helices undergo crossing in the Int2 structure. These key conformational changes between the bent-closed, Int1, Int2, and the extended-open states are captured by the CVs, 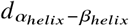 and 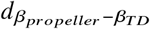.(11, 31) Accordingly, we define the initial path as an equal arc length string parameterized by *N*+1=19 equidistant points at a distance of 0.75 nm in the 2D physical CV space. These 19 points were generated through three linear interpolations, with each interpolation performed between two of the four manually selected locations from the projections of the bent-closed, Int1, Int2, and extended-open states sampled in Tong et al. (18) in the 2D physical CV space (see SI Fig. S1). We label the set of the 19 linearly interpolated points in 2D physical CV space as ***ϕ***_*λ*_ = {*ϕ*_0_, *ϕ*_1_, .., *ϕ*_18_ }, where *λ* identifies each point in the set with *λ* = 0 and *λ* = 18 corresponding to the points in the bent-closed state and extended-open state, respectively.

### Relaxation of the path using the FTS method

We refine our initial estimate of the minimum free energy deadbolt activation pathway using the FTS method with the Voronoi tessellation algorithm(32). This algorithm aims to identify the principal curve connecting two metastable states in the CV space, as defined by the center of a tube that carries the highest flux of transitions between two metastable states. In this work, we use the terms minimum free energy pathway and principal curve interchangeably.

First, we perform a Voronoi tessellation of the 2D physical CV space spanned by 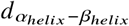 and 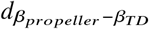 using the 19 ***ϕ***_*λ*_ from the initial guess path as the node centers (Fig. 2b). This partitions the CV space along the path by identifying regions that are closer to a given image 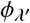 than any other images ***ϕ***_*λ*_, where *λ*^′^ ≠ *λ*. The proximity of two points **z** and **z**^′^ in the CV space can be measured by 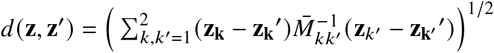. The metric tensor 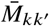 accounts for the curvilinearity of CV space. For the two distance-based CVs, 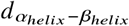 and 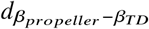, the metric tensor is the identity matrix scaled by an immaterial constant and, therefore, we drop 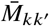 in all future discussion. The boundary Voronoi tessellation regions corresponding to the images *ϕ*_0_ and *ϕ*_18_ represent the bent-closed state and extended-open state, respectively.

Next, the initial guess path was iteratively relaxed by evolving it in the 2D physical CV space spanned by 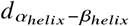 and 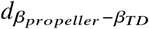 (Fig. 2b). This involves two steps. In the first step, the location of the images are refined according to 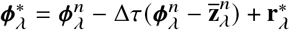, where *n* refers to the iteration number. The images 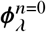 in the iteration *n* = 0 correspond to those in the initial guess path ***ϕ***_*λ*_ defined in the previous subsection, and 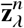 refers to the average location of the system in the Voronoi cell corresponding to the image *λ* in iteration *n*. We estimate 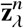 by performing AA MD simulations, which we elaborate in the next subsection. The evolution time step parameter 0 ≤ Δ*τ* ≤ 1 controls the relative weight provided to 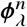 and 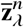 in each iterative update of the path, which we set to Δ*τ* = 0.1. The constraint term 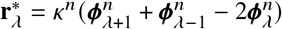 with 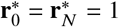 ensures smoothness of the string 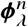 by damping the influence of any statistical fluctuations in 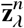 during an update. *k*^*n*^ = *kN*Δ*τ* with *κ*=0.1 refers to an adjustable parameter for controlling the strength of the smoothening, which because of its proportionality to the number of images *N* helps in minimizing the discretization errors or kinks along the string associated with the finite number of *N* used for defining the continuous string in practice. In the second step, a reparameterization step was applied to obtain the images of the path in the next iteration 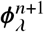. This involves interpolation of a piecewise linear curve through the revised images 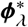. The reparameterization step ensures the equidistant image constraint 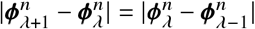 was satisfied for representing the string in its intrinsic geometric parameter form with arc length parameterization defined initially and additionally helps in achieving proper sampling of the system along the string for the estimation of 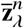.

### Exploration of the configurational space along the path using AA MD simulations

To iteratively refine the initial guess of the minimum free energy deadbolt activation pathway using the FTS method(32), we need to sample the configurational space along the path in each iteration *n* for estimating 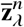. For this, we perform explicit water AA MD simulations of *α*_*IIb*_ *β*_3_ integrin in lipid bilayer using AA integrin structures that correspond to each of the 19 images 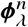 in the 2D physical CV space (Fig. 2c). We construct the AA structures of *α*_*IIb*_ *β*_3_ integrin for sampling in the Voronoi cells corresponding to each *λ* in each iteration *n* by identifying the structures closest to the images 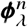 from the samples collected in a previous iteration *n* − 1.

For the first iteration *n* = 0, we construct the AA structures of *α*_*IIb*_ *β*_3_ integrin (SI Fig. S2-SI Fig. S3) by employing the multiscale data-driven framework developed in our prior work.(31). Briefly, this framework combines non-linear manifold learning and deep generative modeling to generate AA *α*_*IIb*_ *β*_3_ integrin structures corresponding to each of the 19 images of the initial string, 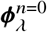. In SI Sec. S1, we provide the details of this approach. To prepare the resulting *α*_*IIb*_ *β*_3_ integrin structures for explicit water AA MD simulations within the lipid bilayer, we added the critical cofactor ions in the binding sites of *α*_*IIb*_ *β*_3_ integrin using the Orient package in VMD 1.9.4(33) to first align the protein with the principal axes and then employing the TopoTools(34) package to insert the ions. In particular, we added one Mg^2+^ ion to the metal ion-dependent adhesion site (MIDAS) in the *β*_*A*_ domain. We added six Ca^2+^ ions to the four binding sites in the *β*_*propellar*_ domain, one synergistic metal binding site (SyMBS), and one adjacent to MIDAS (ADMIDAS) sites in the *β*_*A*_ domain. Residues listed in SI Tab. S1 were used to align each binding site. An X-ray crystallography structure corresponding to the *α*_*IIb*_ *β*_3_ integrin in the bent-closed state downloaded from the Protein Data Bank (PDB: 3ZDY(35)) was used as the reference structure for placement of the two types of cations in all the 19 images.

Following the incorporation of the ions, we determined the spatial positioning and orientation of the 19 AA integrin structures within the lipid bilayer. This was achieved using the position of the protein in membranes (PPM) 3.0 method(36) implemented in the webserver(37) at https://opm.phar.umich.edu/ppm_server3. Briefly, this approach determines the optimal distance (*d*), tilt (*ϕ*) and rotation (*τ*) of a given protein with respect to the membrane normal by minimizing the free energy (Δ*G*_*transf*_ (*d*, *ϕ*, *τ*)) required for transferring the protein from water to a membrane environment.(38) PPM 3.0 approximates Δ*G*_*trans f*_ (*d*, *ϕ*, *τ*) using the universal solvation model(39) for anisotropic environments. To perform these orientation calculations, we uploaded each of the 19 initial AA *α*_*IIb*_ *β*_3_ integrin structures with cofactor ions to the PPM 3.0 webserver and selected the closest available target membrane environment i.e., DOPC (1,2-dioleoyl-sn-glycero-3-phosphocholine) for conducting these orientation and placement calculations. We then passed the optimally oriented *α*_*IIb*_ *β*_3_ integrin structures with cofactor ions to the Membrane Builder tool in CHARMM GUI(40–42) to build the AA lipid bilayer in water at biophysical conditions of 0.15 M NaCl concentration. The lipid bilayers were aligned along the XY dimensions of a rectangular simulation box. Their dimensions were adjusted based on the 3:1 DOPC:DOPS (1,2-dioleoyl-sn-glycero-3-phospho-L-serine) molar ratio in CHARMM GUI with the initial guess of the XY dimensions set to accommodate at least 2 nm of distance between the periodic images of integrin. CHARMM GUI was used for setting the Z dimension of the simulation box based on the thickness of the bulk water from integrin above and below the membrane. SI Tab. S2 contains the details of the initial simulation box size and the system composition comprising each of the 19 AA *α*_*IIb*_ *β*_3_ integrin-lipid bilayer systems along the initial string.

We modeled the AA integrin-lipid bilayer complex using the CHARMM36 protein and lipid force field(43–45). Water molecules were represented by CHARMM compatible TIP3P water model(43, 46, 47). Next, we conducted steepest descent energy minimization to eliminate forces greater than 1000 kJ/mol/nm, followed by six step equilibration cycles for slowly equilibrating the 19 *α*_*IIb*_ *β*_3_ integrin-bilayer complex systems in a NPT ensemble at 310 K and 1 bar pressure.(40–42) Additional details on the energy minimization runs and equilibration procedure along with the simulation parameters can be found in SI Sec. S2.

Finally, we estimated the configurational average of the system, 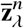, by performing unbiased MD simulations of the 19 AA *α*_*IIb*_ *β*_3_ integrin-lipid bilayer systems corresponding to each image *λ* in each iteration *n* independently by following the conditions for FTS method. Specifically, we employed OpenMM 7.7(48) patched with SEEKR2 library(49) to run the AA MD simulations using a custom Langevin integrator called MmvtLangevinIntegrator(49) that respects the reflecting boundary conditions when the system strikes the Voronoi partitions of each image in the 2D physical CV space. This allowed configurational sampling of each system within their respective Voronoi cells in the 2D physical CV space, thereby providing samples 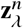 for estimating their average, 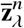. The OpenMM input scripts for performing these AA MD simulations were prepared using CHARMM GUI(40–42) and modified by adding the definitions of the Voronoi cells to use the SEEKR2 library(49). In each iteration and for each image, the MD simulations were conducted for 200 ps by regulating temperature at 310 K using a friction coefficient of 1 ps^−1^ and the integration time step was set to 2 fs. We employed the semi-isotropic Monte Carlo membrane barostat(50) implemented in OpenMM 7.7(48) to regulate the pressure at 1 bar with a frequency of 100 timesteps for the Monte Carlo volume changes because the pressure coupling was applied isotropically along x- and y-axes while the pressure along the z-axis was allowed to vary independently. We constrained the bonds connected to hydrogen atom in integrin and lipid bilayer using the CMMA algorithm(51) in OpenMM 7.7(48).

In each iteration, the 200 ps of AA MD simulations took ∼75 min to ∼140 min depending on the number of atoms in a system corresponding to a image along the string. We allocated 4 GPUs with 40 CPUs to systems containing more than 1.5M atoms. For the remaining systems, we employed 1 GPU with 10 CPUs. The GPUs were either Nvidia RTX 2080 Ti or Titan-V. Altogether, this work consumed approximately 440,000 CPU-h and 40,000 GPU-h of compute time. The OpenMM scripts for running all the simulations with FTS method along with the relaxed AA structures from the relaxed path presented in this work as well as the Jupyter notebooks used for analysis can be accessed at https://github.com/Ferg-Lab/principalcurve_integrin_structures.git.

### Assessment of the convergence of the paths

We periodically assessed the convergence of the relaxation of the FTS method by monitoring the change in the distance between the evolved paths with iterations (Fig. 2d). We employed the discrete Fréchet distance(52, 53) to measure the distance between the paths and their relaxation to the minimum free energy pathway. The discrete Fréchet distance measures the minimum distance between two given discretized curves by traversing both the curves in the same direction without backtracking.(53, 54) In other words, it captures the similarity of the shape of two discrete curves when arranged in a specific order. In our context, it measures the similarity between two strings traversed from the bent-closed state to extended-open state. We employed the Python library frechetdist 0.6(55) for computing the discrete Fréchet distances in this work.

We consider the paths evolved by the FTS method to be converged when the distance between the string 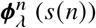 in iteration *n* and the string 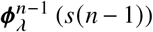 in the preceding iteration *n* − 1 reaches a plateau in both the Fréchet distance and mean distance. Additionally, we monitor the convergence of the distance between *s* (*n*) and the curve formed by the configurational average 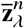 in an iteration *n* (*r* (*n*)). This distance between *s* (*n*) and *r* (*n*) tracks the evolution of the string towards the principal curve(32). Accordingly, we terminate the FTS method cycle when this distance starts to plateau with iterations.

We iteratively refined the initial guess for the minimum free energy deadbolt activation pathway for a total of 396 iterations to achieve convergence of the paths evolved using the FTS method. While the path converged in the first ∼ 50 iterations, we observed erroneous transition structures especially in the tail membrane helices region (SI Fig. S4). We discovered that these unphysical transition structures were a consequence of the incorrectly resolved tail membrane regions in the integrin structures along the initial guess pathway (SI Fig. S2-SI Fig. S3). To address this issue, we re-generated the 19 initial AA integrin structures using our initial structure generation approach discussed in SI Sec. S1 along the relaxed path. We found that these structures displayed physically plausible transitions including the tail membrane helices region along the relaxed path (SI Fig. S5-Fig. S6) corresponding to iteration 249. Subsequently, we continued the FTS cycle until the paths with these revised structures converged, which took an additional 146 iterations resulting in a total of 396 iterations for the overall FTS relaxation process.

### Preparation of partial or full-length experimental structures for analysis

In order to relate the minimum free energy activation pathway determined using the FTS method, we sought to compare our predictions of the structures along the relaxed path to the available experimentally resolved structures of *α*_*IIb*_ *β*_3_ integrin in various intermediate states. For this, we identified and downloaded 68 experimentally resolved structures of the *α*_*IIb*_ *β*_3_ integrin from the PDB. The resulting structures include entries that were resolved using X-ray crystallography, NMR spectroscopy, and cryo-EM methods. SI Tab. S3-SI Tab. S5 contains a brief summary of these PDB entries. To remove non-integrin entities, add terminal patches and disulfide bonds, and separate files containing multiple structures for analysis in the X-ray crystallographic and NMR PDB files, we employed the CHARMM GUI PDB Reader & Manipulator tools(40, 56). This step was not necessary for the cryo-EM PDB entires. The C_*α*_ atoms in the cleaned structures were then aligned with the relaxed structures from the last iteration in the FTS cycle for analysis. For performing this structure alignment, we identified all the common residues between the experimental and predicted structure from FTS calculations using a custom VMD(33) tcl script.

## RESULTS AND DISCUSSION

Our goal in this work is to resolve the full-length AA structures of *α*_*IIb*_ *β*_3_ integrin along the minimum free energy pathway connecting the bent-closed state and extended-open state. We achieve this by employing the FTS method to relax an initial guess for the deadbolt activation pathway generated using our multiscale generative modeling approach detailed in Dasetty et al. (31). Below, we first demonstrate the relaxation of the path to the minimum free energy activation pathway using the FTS method. Then, we assess the validity of the relaxed path by comparing the structures along it with the experimentally resolved structures. We follow this with a discussion on the structural changes along the relaxed pathway to provide insights into the activation mechanism of *α*_*IIb*_ *β*_3_ integrin.

### Relaxation of the initial guess for the deadbolt activation pathway using the FTS method

We determine the minimum free energy activation pathway between the bent-closed state and the extended-open state of *α*_*IIb*_ *β*_3_ integrin using the FTS method(32) in a 2D physical CV space spanned by 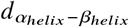 and 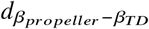 (SI Fig. S1). These CVs are selected because they capture the key conformational transitions between the bent-closed state and the extended-open state.(11, 31) We seed the FTS method with an initial guess pathway that traces the deadbolt activation mechanism within the CV space constructed using our previously reported multi-scale generative approach(31). We then iteratively refine the path using the FTS method with Voronoi tessellation(32), where we run AA MD simulations to relax the configurations comprising the path to the local free energy minimum pathway. The evolution of paths in the FTS method procedure is monitored by tracking the changes in the location of images in the 2D physical CV space. Fig. 3(a) shows the relaxation of the path towards the minimum free energy path as a function of the number of iterations in the FTS method. Additionally, Fig. 3(a) presents the projection of 2450 AA MD configurations in the CV space for reference on the location of the known full-length conformational states of *α*_*IIb*_ *β*_3_ integrin. These reference configurations are sampled by Tong et al. (18) using four experimentally determined full-length conformations(9) (Fig. 3(a)) of the *α*_*IIb*_ *β*_3_ integrin, which correspond to the bent-closed state, two extended intermediates (Int1, Int2), and the extended-open states, respectively. Fig. 3(a) shows there are no transitions between these four conformational states within the CV space during the 500 ns AA MD simulations conducted in the previous work(18), illustrating the challenge in sampling the transitions between these states using conventional unbiased AA MD simulations.

**Figure 3:**
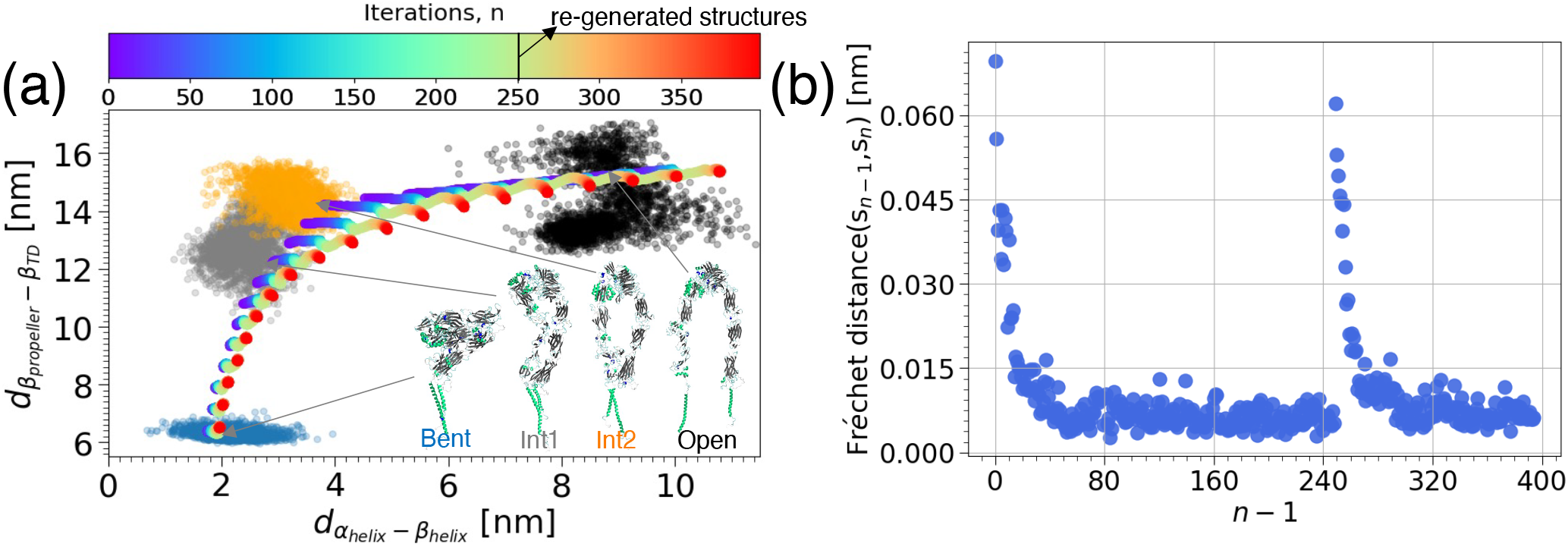
Relaxation of the initial guess for the deadbolt activation pathway using the FTS method. (a) Evolution of the paths by the FTS method procedure in the 2D physical CV space spanned by 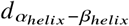 (x-axis) and 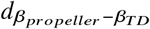 (y-axis). The path in each iteration is constituted by 19 equidistant images, 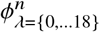. The color scale from violet to red correspond to changes in the path with iterations during the FTS method relaxation. The projection of 2450 MD AA configurations sampled in prior work by Tong et al. (18) with initial structures(9) corresponding to the bent-closed state (translucent blue), two extended intermediate states (Int1: translucent gray, Int2: orange), and the extended-open (translucent black) states are shown for reference of the known full-length structures of *α*_*IIb*_ *β*_3_ integrin in the CV space. Visual snapshots of the configurations corresponding to the four states are rendered using VMD(33). (b) Fréchet distance between the string in iteration *n* (*s*_*n*_) and iteration *n* − 1 (*s*_*n*−1_). The Fréchet distance (*s*_*n*−1_, *s*_*n*_) plateaus at about iteration 50 indicating the convergence of the string with iterations. In iteration 249, we have re-generated the 19 AA structures to correct the unphysical transitions in the tail membrane helices followed by an additional 146 iterations resulting in a total of 396 iterations for the overall FTS relaxation.

In Fig. 3(b), we quantify the convergence of the FTS with successive iterations using the Fréchet distance as the path similarity metric. We have observed that the Fréchet distance between the path in an iteration *n* (*s* − (*n*) ) and the iteration *n* − 1 (*s* (*n* −1) ) decreases rapidly and reaches a plateau after iteration ∼ 50 (Fig. 3(b)), indicating the convergence of the string. However, we observed erroneous transition structures (SI Fig. S4), particularly in the tail membrane helices region. Specifically, we observed an invalid structure (SI Fig. S2-SI Fig. S4) corresponding to the image *λ* = 11, which is inconsistent topologically with its neighbors due to different crossing of tails. This unphysical transition structure is a consequence of the incorrectly resolved tail membrane helices in the integrin structures along the initial guess pathway. To address this issue, we re-generated all the 19 initial AA integrin structures along the converged path at iteration 249 to generate an ensemble of structures along the string that did not suffer from this topological incompatibility and continued the FTS cycle to further relax the path with the revised structures. SI Fig. S5-SI Fig. S6 shows that the transitions in the revised initial structures are gradual, including the separation of the tail membrane helices along the path. This re-setting of the initial integrin structures initially triggered an abrupt increase in the Fréchet distance between *s* (*n*) and *s* (*n* − 1) at iteration 250 depicted in Fig. 3(b). In approximately 50 iterations after this reset that corresponds to the iteration ∼ 300, the Fréchet distance *s* (*n* ) and *s* (*n* −1) experienced a second decline and plateaued, indicating convergence.

We postulate that the rapid convergence of the path implies that the initial guess deadbolt activation pathway was a good approximation for the *α*_*IIb*_ *β*_3_ integrin activation process but it is also possible that the string is trapped in a local optimum and has failed to identify the true minimum free energy path. To answer this and understand the importance of the relaxed path, we now proceed to compare the structures along the relaxed path (SI Fig. S7) with structures resolved by a variety of experimental methods.

### Comparison of the structures along the relaxed path with experimentally resolved structures

The images along the relaxed path from the FTS method follow a typical deadbolt like mechanism model, wherein first 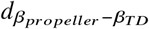 increases gradually and then the tail helices separate as reflected in the increase in 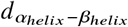 (Fig. 4(a)). However, this minimum free energy deadbolt activation path may not be the true minimum free energy activation path as it is possible that the pathway is trapped in a local free energy minimum during FTS relaxation. To evaluate the physical plausibility of the terminal FTS path we compare predicted structures along the path with 68 experimentally resolved structures of *α*_*IIb*_ *β*_3_ integrin collected using X-ray crystallography, NMR spectroscopy, and cryo-EM. In Fig. 4(b), we present this comparison by reporting the RMSD of each of the 68 experimental structures to its closest predicted structure along the terminal FTS string.

**Figure 4:**
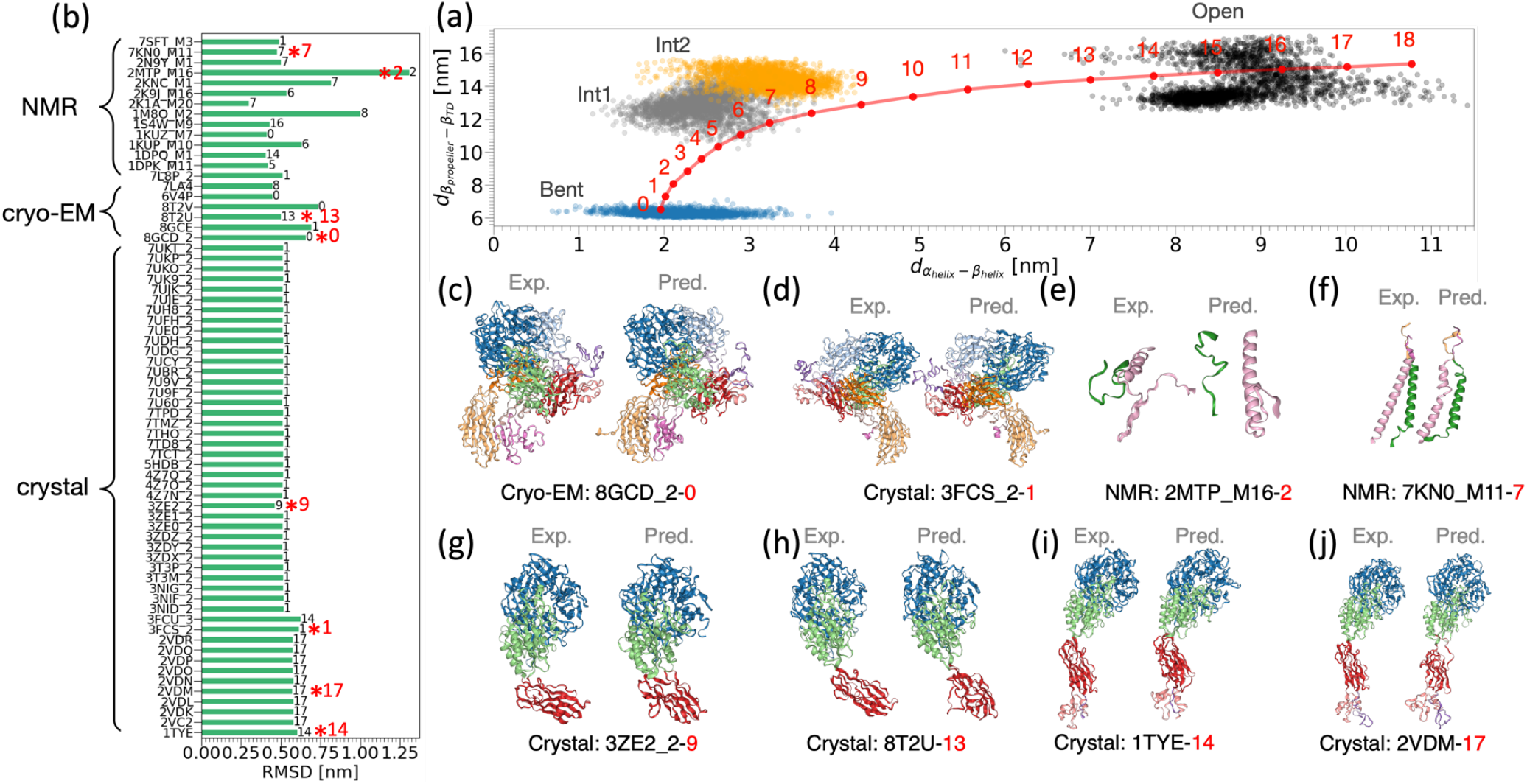
Comparison of the *α*_*IIb*_ *β*_3_ integrin structures along the relaxed pathway with experimentally resolved structures. (a) Location of the images (solid red markers) from the relaxed path (solid red line) in the 2D physical CV space spanned by 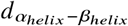 (x-axis) and 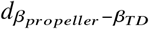 (y-axis). The translucent blue, gray, orange, and black markers correspond to the projection of the 2450 full-length AA MD configurations sampled in Tong et al. (18) with initial structures from cryo-electron microscopy(9) in bent-closed, Int1, Int2, and extended-open states in the CV space, respectively. (b) RMSD between the structures from the relaxed pathway and 68 structures resolved using X-ray crystallography, NMR spectroscopy, and cryo-EM techniques. The PDB IDs of the experimentally resolved structures are shown along y-axis and their RMSD values compared to a structure in the relaxed pathway are shown along x-axis. The image number *λ* shown on the bar indicate the structure in the relaxed pathway with the minimum RMSD from the experimentally resolved structure. (c)-(j) Visual comparisons of selected experimental structures (left) and their corresponding closest structure in the relaxed pathway (right). The PDB IDs of the experimentally resolved structures and the location of the structure along the relaxed pathway in the CV space shown in (a) are provided below the snapshots. The corresponding RMSD values are highlighted using red asterisk in (b) along with the image labels in red. NGLview 3.0.3(57) is used for rendering the structures and colors used to identify the various domains labeled in Fig. 1.

All the structures resolved using X-ray crystallography and cryo-EM methods have a RMSD of less than 1 nm with at least one of the structures corresponding to an image along the relaxed string. These experimental structures include partially and fully resolved ectodomains of *α*_*IIb*_ *β*_3_ integrin, which are considered to be transient structures along the activation process (SI Tab. S3-SI Tab. S4). Two of the cryo-EM structures 8GCD and 8GCE were resolved after the publication of our prior work(31) and include the complete ectodomain along with a predicted transmembrane helices orientation.(58) These new structures 8GCD and 8GCE are also found to be in good agreement to the relaxed structures corresponding to images *λ* = 0 and *λ* = 1 of the relaxed path (Fig. 4(a)) near the bent-closed state with a RMSD of 0.768 nm and 0.695 nm, respectively. If we exclude the transmembrane helices that are not well-resolved experimentally, the RMSD of 8GCD (8GCD_2 in Fig. 4(b)) with structure corresponding to *λ* = 0 further reduces to 0.657 nm.

All the available NMR structures contain only the transmembrane helices (SI Tab. S5). Given the ensemble of structures resolved from NMR experiments, we have isolated one NMR structure from each PDB deposit that has the minimum RMSD with a given image along the relaxed path and shown in Fig. 4(b). Even these transmembrane helices and cytoplasmic domains resolved using NMR spectroscopy have a good match (RMSD of 0.299-1.309 nm) with one of the structures along the relaxed path (Fig. 4(a)).

Example all-atom structures comparing the experimentally resolved structure with a relaxed structure along the relaxed path are shown in Fig. 4(c)-(j). These structures are arranged according to the location of the images with a minimum RMSD in the 2D physical CV space (Fig. 4(a)). This visual analysis again shows that our computationally resolved structures are in good agreement with the experimentally resolved structures. In addition, they can potentially help in identifying the location of the experimental structures in the 2D physical CV space especially as some of these structures are not full-length and don’t contain the domain pairs corresponding to the x- and y-axes.

To identify if the close agreement in the structures is because of the FTS relaxation procedure, we additionally compared the structures generated along the initial guess deadbolt activation pathway with the experimentally resolved structures. SI Fig. S8 shows that the RMSD of even the structures along the initial pathway with the experimentally resolved structures is generally better than 1 nm. Given that the initial structures are predictions themselves based on our multi-scale data-driven framework(31), the agreement of the structures along the initial guess pathway with experiments lends confidence to the initial structures employed to seed the FTS method. Furthermore, it suggests that the FTS relaxation procedure primarily relaxed their position in the 2D physical CV space while preserving the experimentally consistent features of the initial structures.

Taken together, the close correspondence of the predicted structures along the relaxed FTS pathway with experimental structures provides supporting evidence for the physical plausibility of the predicted activation pathway and the capacity of our computational approach to predict physically plausible transient structures of *α*_*IIb*_ *β*_3_ integrin, which are otherwise challenging to resolve at full-length and AA resolution by both experiment and conventional computational methods. To demonstrate its application in elucidating the activation mechanism of *α*_*IIb*_ *β*_3_ integrin, we will next analyze the gross conformational changes in these structures along the relaxed pathway.

### Changes in the full-length *α*_*IIb*_ *β*_3_ integrin structure along the relaxed path

We now proceed to interpret the structural changes along it to mechanistically understand the structural transition from the bent-closed state to the extended-open state. In Fig. 5(a), we show the changes in the full-length structures along the minimum free energy pathway between the bent-closed state and extended-open state of *α*_*IIb*_ *β*_3_ integrin. The separation between the *β*_*propellar*_ and the *β*_*TD*_ of the *α*_*IIb*_ *β*_3_ integrin is gradual along the relaxed string (Fig. 5(b)). In contrast, the transmembrane helices maintain a conformation similar to the bent-closed state up to the image *λ* = 9, after which they gradually separate to the one in the extended-open state (Fig. 5(b)). This trend is additionally followed by the hybrid domain swing-out with an angle similar to bent-closed state up to the image *λ* = 9, with image 9 through 11 representing a transition state, and the successive images representing an extended-open state. Additionally, the opening of the hybrid domain when comparing PDB: 3T3P (bent-closed state)(59) to PDB: 2VDR (extended-open state)(60) is ∼ 33^°^, which is in good agreement with our measured values ranging from ∼ 0^°^ to ∼ 36^°^ as shown in SI Fig. S9.

**Figure 5:**
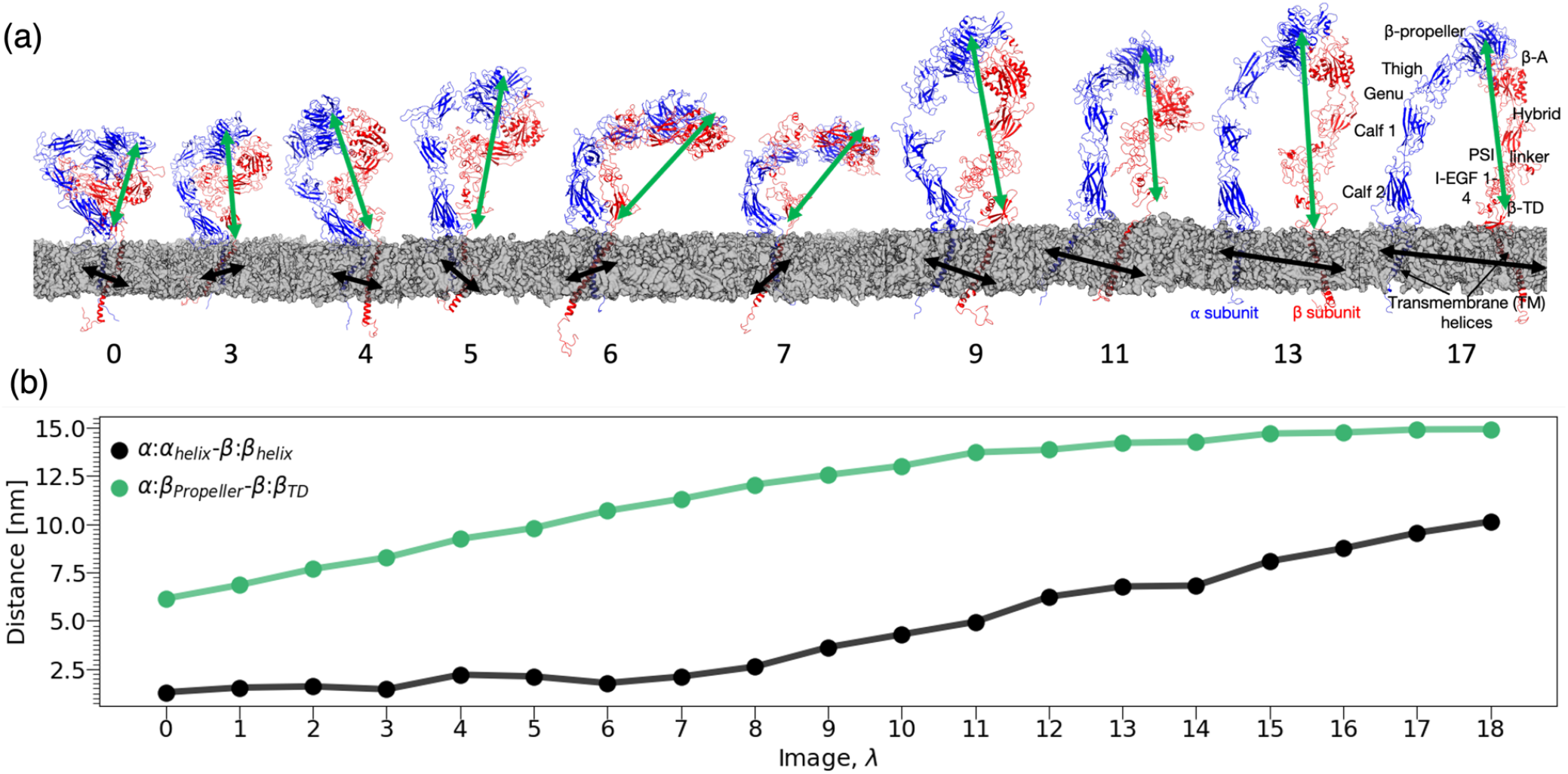
Changes in the full-length structure of *α*_*IIb*_ *β*_3_ integrin embedded in lipid bilayer along the relaxed path, which represents the minimum free energy pathway between the bent-closed state and extended-open state. (a) Visualizations of *α*_*IIb*_ *β*_3_ integrin with the *α* and *β* subunits colored in blue and red, respectively. The image numbers along the string are shown below the snapshots of the structures. Lipid bilayers are illustrated using transparent gray surface representation in Chimera(13, 14). Water molecules and ions are not shown for clarity. Green and black double ended arrows are used to highlight the changes in the 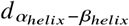 and 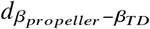, respectively along the relaxed pathway. (b) Changes in 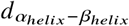 and 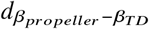 along the relaxed pathway between the bent-closed state and extended-open state are shown using green and black colors, respectively.

The changes in the structure following the deadbolt mechanism model in Fig. 5(a) also reveal potential changes in the different domains and the orientation of the *α*_*IIb*_ *β*_3_ integrin with respect to the lipid bilayer during the activation process. For example, SI Fig. S10-SI Fig. S12 reveal that the change in 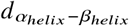 along the string is strongly correlated with changes in distance between several other domain pairs such as thigh and E3 domain, thigh and E4 domain, and Calf-1 and E4 domain along the string. Similarly, the change in 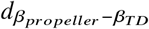 along the string is strongly correlated with the changes in the distance between domain pairs such as *β*_*propellar*_ and Calf-1, *β*_*propellar*_ and Calf-2, and *α*_*helix*_ and *β*_*A*_ along the string. In addition to these concerted domain rearrangements, the ectodomain of integrin undergoes orientation changes with respect to the lipid bilayer as observed in Fig. 5(a). In SI Fig. S13, we quantify the orientation transitions by presenting the variations of *θ*-***ϕ*** along the string, where *θ* and ***ϕ***, respectively, correspond to the polar and azimuthal angles of the COM of subdomains in the two subunits with respect to the lipid bilayer normal. This plot reveals that the changes in *θ* of both the *β*_*propellar*_ and *β*_*A*_ domains decrease and are correlated with a Pearson correlation coefficient of *ρ*=0.94 along the relaxed path, which captures the extension of the headpiece during the transition. The azimuthal angle, ***ϕ***, of *β*_*propellar*_ and *β*_*A*_ are correlated with a *ρ*=0.75 and undergo only minor changes, but with larger fluctuations in *β*_*propellar*_ domain, indicating their coordinated fluctuations along the lipid bilayer during their transition from the bent-closed to the extended-open state.

Collectively, the above results indicate that the transition from the bent-closed to the extended-open state through the minimum free energy pathway involve concerted rigid domain movements along with fluctuations in the orientation of headpiece with the lipid bilayer.(58) These concerted rearrangement of domains during the activation dynamics could be critical for integrin to act as a mechanotransducer in various cellular responses.(21)

## SUMMARY AND CONCLUDING REMARKS

Understanding the conformational activation of the integrin heterodimer is of fundamental importance in elucidating its cellular functions and in designing integrin targeting therapeutics. However, resolving the conformational changes along the activation pathway between the bent-closed state and the extended-open state remains challenging for both experimental and computational methods. In this work, we use the FTS method to identify the minimum free energy activation pathway and the corresponding AA structures along it. We seed the FTS method by constructing an initial string using the deadbolt activation model in a 2D physical CV space spanned by 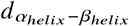 and 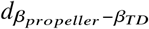 using our multiscale data-driven framework(31) to generate physically plausible AA structures for the images along the initial guess pathway. The final relaxed pathway from the FTS cycle are in good agreement with 68 experimentally resolved (partial) structures of *α*_*IIb*_ *β*_3_ integrin. Additionally, the structures revealed several correlated changes between the distances of the domains of integrin along with variations in the orientation of the ectodomain headpiece with respect to the lipid bilayer. These insights and the predicted transient structures have potential applications in designing therapeutics to treat integrin dysfunction or malfunction.(61)

One of the limitations of this work is that the identified minimum free energy pathway is a function of the choice of the initial string especially because of potential limited sampling in each Voronoi cell along the string due to the large size and slowly evolving nature of the integrin system. Therefore, the relaxed pathway corresponds to the potential deadbolt mechanism model that served as the basis for our initial string. The quick convergence of the string with iterations suggests that the initial guess for the string generated by our multi-scale generative modeling approach(31) presented a good initial guess for the deadbolt activation pathway and the good agreement with structures along the relaxed string with experimentally resolved (partial) structures lend physical plausibility to the predicted activation pathway. In future work, we would like to apply similar approaches to estimate relaxed activation pathways commencing from initial guesses based on the switchblade and concerted activation mechanisms and identify which predicted pathways show the best agreement with experimental data. Moreover, we would like to apply path CV enhanced sampling methods(24, 62) to resolve the free energy profile along the relaxed strings to identify metastable states as putative thermodynamic targets. Finally, we would like to extend this approach to other integrin heterodimers and, more generally, other biomolecular systems for which structural transitions are rare events and challenging to sample by conventional techniques.

## AUTHOR CONTRIBUTIONS

SD and ALF designed the research. SD and REC performed the simulations. All authors analyzed the data and wrote the manuscript.

## ACKNOWLEDGMENTS

This work was supported by the U.S. Department of Energy, Office of Science, Basic Energy Sciences, under Award #DE-SC0023318. We gratefully acknowledge computing time provided by the University of Chicago Research Computing Center (https://rcc.uchicago.edu), the University of Chicago high-performance GPU-based cyberinfrastructure supported by the National Science Foundation under Grant No. DMR-1828629. We thank Dr. Gregory A. Voth for helpful discussions and insights.

## SUPPLEMENTARY MATERIAL

Supporting material can be found online.

