## Supplementary Information for "Prediction of *α*_*IIb*_*β*_3_ integrin structures along its minimum free energy activation pathway"

### S1 Generation of the initial AA structures corresponding to the images of the initial guess pathway

We estimated the configurational average of the system within each Voronoi cell during each iteration $n$ of the FTS method using AA MD simulations. To conduct these AA MD simulations, we employed initial AA structures as those closest to the image locations in the 2D physical CV space derived from AA MD samples collected in a previous iteration. For the initial guess pathway, we generated the initial AA structures by utilizing our multiscale data-driven framework.[S1] This method integrates non-linear manifold learning and deep generative modeling methods to learn and generate the transient structures of $\alpha_{IIb}\beta_3$ integrin. In brief, we first learned a unified low dimensional representation of bent-closed, Int1, Int2 and the extended-open conformational states of $\alpha_{IIb}\beta_3$ integrin at 300 bead CG resolution using adaptive diffusion maps.[S2] Then, we trained a conditional Wasserstein generative adversarial network[S3–S5] (cWGAN) model to learn the inverse mapping between the unified low-dimensional embedding and the high dimensional $\alpha_{IIb}\beta_3$ integrin structures at 300 bead CG resolution. The trained cWGAN model can generate $\alpha_{IIb}\beta_3$ integrin structures at 300 bead CG resolution at any location within or between the bent-closed, Int1, Int2, and extended-open conformational states in the unified low-dimensional embedding. In Dasetty et al.[S1], we applied this approach to generate 90,000 transient configurations at 300 bead CG resolution to present plausible activation structures between the bent-closed and extended-open conformational states of $\alpha_{IIb}\beta_3$ integrin. In this work, we utilized these transient structures to identify the structures that were closest to the constructed initial images $\boldsymbol{\phi}_{\lambda=\{0,...,18\}}$ in the 2D physical CV space (SI Fig. S1). Snapshots of these structures were depicted in SI Fig. S2.

We upgraded the 300 bead CG configurations to an AA resolution by using targeted MD simulations[S6] (TMD) with the same CG to AA mapping procedure described in our prior work.[S1] This method involves the application of a moving harmonic biasing potential

($V = (\kappa(t)/2)(\text{RMSD} - \text{RMSD}_{\text{center}}(t))^2$) to minimize the RMSD between a reference AA configuration and the CG target structure.[S6,S7] $\kappa(t)$ and $\text{RMSD}_{\text{center}}(t)$ correspond to the time varying harmonic force constant and center of the harmonic potential, respectively. We used three reference AA configurations that correspond to relaxed full length experimental structures in bent-closed, Int1, and Int2 states for the 300 bead CG target structures at images $\boldsymbol{\phi}_{\lambda=0,..,2}$, $\boldsymbol{\phi}_{\lambda=3,...,10}$, and $\boldsymbol{\phi}_{\lambda=11,...,19}$, respectively. Given the missing residues in the AA structure of extended-open state[S8] employed for AA MD simulations in Tong et al.[S9], we opted to employ the AA structure in the Int2 state rather than extended-open state as the reference AA configuration for the image structures ($\boldsymbol{\phi}_{\lambda=11,...,19}$) near the extended-open state. For all images, we linearly scaled the harmonic bias force constant $\kappa(t)$ from 0 to 15000 kJ/mol/nm$^2$ while varying the $\text{RMSD}_{\text{center}}(t)$ from the initial RMSD to $\sim$0 nm in a simulation time of 2.5 ns.

We accelerated all the backmapping TMD simulations by employing the generalized Born implicit solvent model[S10] with the solute and solvent dielectric constants set to 1 and 78.5, respectively. CHARMM36[S11] force field was employed to model the AA integrin with length of all bonds involving a hydrogen atom constrained using the constant constraint matrix approximation (CCMA)[S12] algorithm implemented in OpenMM 7.7.[S13] We conducted these simulations at 310 K using Langevin equations of motion by setting the friction coefficient to 0.1 ps$^{-1}$ and the integration time step to 2 fs. The forces from the moving harmonic bias potential were added to the system by using PLUMED 2.7.3[S14] enhanced sampling software library. All the AA reference configurations were energy minimized until convergence with a tolerance criteria of 10 kJ/mol for the energy of the system prior to initiating the TMD simulations. Snapshots of the resulting backmapped AA structures corresponding to the 300 bead CG configurations shown in SI Fig. S2 are illustrated in SI Fig. S3.

### S2 Energy minimization and equilibration of the initial AA integrin-lipid bilayer complex structures corresponding to the images of the initial guess pathway

We subjected each of the AA integrin-lipid bilayer complex systems corresponding to the 19 initial images to an energy minimization run for a maximum of 5000 steps or until convergence with a tolerance of 1000 kJ/mol/nm for the maximum force in the system. We followed this by the CHARMM GUI[S15–S17] suggested six step equilibration cycles for slowly equilibrating the lipid bilayer and integrin in a NPT ensemble at 310 K and 1 bar pressure. In these six steps, the position restraints applied to all the non-hydrogen lipid atoms in the system were gradually weakened to stabilize the lipid bilayer. Temperature and pressure were regulated by Berendsen thermostat[S18] with 1 ps coupling constant and a semi-isotropic Berendsen barostat[S18] with 5 ps coupling constant and 4.5 $\times 10^{-5}$ bar$^{-1}$ compressibility, respectively in the equilibration runs. The first two equilibration cycles were performed in a NVT ensemble for 125000 steps with 1 fs timestep with the position restraints on the lipid atoms weakened in the second cycle. In the third cycle and fourth, the system was equilibrated in a NPT ensemble for 125000 steps with 1 fs and 2 fs time step, respectively, with the position restraints on the lipid atoms further weakened. The last two cycles were also performed in a NPT ensemble for 125000 steps with 2 fs timestep but with the position restraints on the lipid atoms gradually removed completely. For both energy minimization and the six step equilibration cycle, we employed GROMACS 2019.3.[S19,S20] We used the CHARMM36 force field[S17] suggested settings for the switching the Lennard Jones forces from 1.0. nm to 1.2 nm with the force switch scheme implemented in GROMACS 2019.3.[S19,S20] A short range Coulomb cutoff of 1.2 nm was employed with the long range electrostatics computed using particle mesh Ewald method.[S21] All the bonds with hydrogen atoms were constrained using LINCS[S22] algorithm and the neighbor lists were updated every 20 time steps using Verlet algorithm[S23] during the equilibration cycles. We maintained position restraints to all non-hydrogen atoms of integrin and also the co-factor ions to keep the structures close to the image centers of the Voronoi cells during these equilibration runs.

Table S1: Residues used to position the divalent ions within the AA $\alpha_{IIb}\beta_3$ integrin structure during the initial AA MD simulation setup. We added a single ${Mg_2}^+$ ion to the metal ion-dependent adhesion site (MIDAS) site in the $\beta_A$ domain. Four of the six ${Ca_2}^+$ ions were added to the four binding sites (Site 1, Site 2, Site 3, Site 4) in the $\beta_{propeller}$ domain. The remaining two ${Ca_2}^+$ ions were added to the synergistic metal binding site (SyMBS) and the adjacent to MIDAS (ADMIDAS) sites, respectively in the $\beta_A$ domain.

| Ion | Domain | Binding site | Residues |
|---|---|---|---|
| ${Ca_2}^+$ | $\beta_{propeller}$ | Site 1 | 434, 432, 426, 428, 430 |
| ${Ca_2}^+$ | $\beta_{propeller}$ | Site 2 | 371, 365, 367, 369, 373 |
| ${Ca_2}^+$ | $\beta_{propeller}$ | Site 3 | 299, 297, 303, 305, 301 |
| ${Ca_2}^+$ | $\beta_{propeller}$ | Site 4 | 245, 250, 247, 252, 243 |
| ${Ca_2}^+$ | $\beta_A$ | SyMBS | 219, 217, 158, 220, 215 |
| ${Ca_2}^+$ | $\beta_A$ | AMIDAS | 127, 126, 123 |
| ${Mg_2}^+$ | $\beta_A$ | MIDAS | 220, 121 |

Table S2: Details of the AA MD simulation system corresponding to each image $\lambda$ along the string during the FTS method relaxation procedure.

| $\lambda$ | Integrin | Water | Ions | Lipid | Magnesium | Calcium | Total | Box:X [nm] | Box:Y [nm] | Box:Z [nm] |
|---|---|---|---|---|---|---|---|---|---|---|
| 0 | 26906 | 1043097 | 2384 | 222360 | 1 | 6 | 1294754 | 24.036 | 24.036 | 23.671 |
| 1 | 26906 | 1039635 | 2374 | 221270 | 1 | 6 | 1290192 | 24.017 | 24.017 | 23.601 |
| 2 | 26905 | 1129374 | 2544 | 221815 | 1 | 6 | 1380645 | 24.035 | 24.035 | 25.163 |
| 3 | 26907 | 1149660 | 2581 | 221270 | 1 | 6 | 1400425 | 24.031 | 24.031 | 25.566 |
| 4 | 26906 | 1149561 | 3176 | 214932 | 1 | 6 | 1394582 | 24.021 | 24.021 | 25.600 |
| 5 | 26905 | 1224279 | 2731 | 221270 | 1 | 6 | 1475192 | 24.017 | 24.017 | 26.992 |
| 6 | 26907 | 1115571 | 2529 | 222360 | 1 | 6 | 1367374 | 24.040 | 24.040 | 25.024 |
| 7 | 26906 | 1129638 | 2546 | 221270 | 1 | 6 | 1380367 | 24.009 | 24.009 | 25.293 |
| 8 | 26905 | 1105704 | 2504 | 221815 | 1 | 6 | 1356935 | 24.026 | 24.026 | 24.818 |
| 9 | 26905 | 1314225 | 2888 | 221815 | 1 | 6 | 1565840 | 24.026 | 24.026 | 28.509 |
| 10 | 26906 | 1321974 | 2907 | 221815 | 1 | 6 | 1573609 | 24.037 | 24.037 | 28.677 |
| 11 | 26922 | 1232607 | 2741 | 221815 | 1 | 6 | 1484092 | 24.026 | 24.026 | 27.124 |
| 12 | 26923 | 1438569 | 3119 | 221270 | 1 | 6 | 1689888 | 24.024 | 24.024 | 30.768 |
| 13 | 26925 | 1397865 | 3039 | 221270 | 1 | 6 | 1649106 | 24.027 | 24.027 | 30.025 |
| 14 | 26923 | 1451490 | 3148 | 221815 | 1 | 6 | 1703383 | 24.026 | 24.026 | 31.054 |
| 15 | 26923 | 1406400 | 3057 | 221270 | 1 | 6 | 1657657 | 24.040 | 24.040 | 30.143 |
| 16 | 26922 | 1450362 | 3141 | 221815 | 1 | 6 | 1702247 | 24.031 | 24.031 | 30.951 |
| 17 | 26922 | 1437492 | 3116 | 221270 | 1 | 6 | 1688807 | 24.022 | 24.022 | 30.737 |
| 18 | 26923 | 1414926 | 3075 | 221270 | 1 | 6 | 1666201 | 24.029 | 24.029 | 30.345 |

Table S3: Overview of $\alpha_{IIb}\beta_3$ integrin structures resolved using X-ray crystallography.

| PDB ID[1] | Description | State | Resolved domains | Reference |
|---|---|---|---|---|
| 1TYE(1/2/3) | $\alpha_{IIb}\beta_3$ headpiece bound to cacodylate ion. | Extended-open state with high affinity $\beta_A$ domain. | $\beta_{Propeller}$, $\beta_A$, Hybrid, PSI | Xiao et al.[S24] |
| 2VDK(1), 2VDL(1), 2VDM(1), 2VDN(1), 2VC2(1), 2VDO(1), 2VDP(1), 2VDQ(1), 2VDR(1) | $\alpha_{IIb}\beta_3$ headpiece bound to: none, none, tirofiban, eptifibatide, L-739758 HHLGGAKQAGDV, LGGAKQAGDV, HHLGGAKQRGDV, LGGAKQRGDV, respectively. | Extended-open state with high affinity $\beta_A$ domain. | $\beta_{Propeller}$, $\beta_A$, Hybrid, PSI, I-EGF domain 1 | Springer et al.[S25] |
| 3FCS(1/2) | Ectodomain of $\alpha_{IIb}\beta_3$ with no ligand. | Bent-closed state with low-affinity $\beta_A$ domain. | $\beta_{Propeller}$, Thigh, Calf-1, Calf-2 , $\beta_A$, hybrid, PSI, I-EGF 1-4, $\beta_{tail}$. | Zhu et al.[S26] |
| 3FCU(1/2/3) | $\alpha_{IIb}\beta_3$ headpiece with no ligand. | Extended-open state with high affinity $\beta_A$ domain. | $\beta_{Propeller}$, $\beta_A$, Hybrid, PSI | Zhu et al.[S26] |
| 3NID(1/2), 3NIG(1/2), 3NIF(1/2) | $\alpha_{IIb}\beta_3$ headpiece bound to none, none, RUC-1, respectively. | Bent-closed state. | $\beta_{Propeller}$, $\beta_A$, Hybrid, PSI, I-EGF 1. | Zhu et al.[S27] |
| 3T3M(1/2), 3T3P(1/2) | $\alpha_{IIb}\beta_3$ headpiece bound to: RUC-2, none, respectively. | Bent-closed state. | $\beta_{Propeller}$, $\beta_A$, Hybrid, PSI. | Zhu et al.[S28] |
| 3ZDX(1/2), 3ZDY(1/2), 3ZDZ(1/2), 3ZE0(1/2), 3ZE1(1/2), 3ZE2(1/2) | $\alpha_{IIb}\beta_3$ headpiece bound to: GRGDSP with varied $Mn^{2+}/Ca^{2+}$ and $Mg^{2+}/Ca^{2+}$ concentrations. | Bent-closed/Bent-closed, Intermediate/Bent-closed, Intermediate/Intermediate, Intermediate/Intermediate, Intermediate/Intermediate, Intermediate/Extended-Open. | $\beta_{Propeller}$, $\beta_A$, Hybrid, PSI. | Zhu et al.[S29] |

Continued on next page

[1]Numbers within braces refers to the model number present in the PDB structure file. For example, (1/2) refers to the indices of the two models resolved using X-ray crystallography.

Table S3: Overview of $\alpha_{IIb}\beta_3$ integrin structures resolved using X-ray crystallography.

| PDB ID | Description | State | Resolved domains | Reference |
|---|---|---|---|---|
| 4Z7N(1/2), 4Z7O(1/2), 4Z7Q(1/2), 5HDB(1/2) | $\alpha_{IIb}\beta_3$ headpiece bound to: AGDV. | 5 intermediate states. | $\beta_{Propeller}$, $\beta_A$, Hybrid, PSI. | Lin et al. [S30] |
| 7L8P(1/2), 7UJK(1/2), 7UE0(1/2), 7UCY(1/2), 7TCT(1/2), 7UJE(1/2), 7TMZ(1/2), 7U9F(1/2), 7U9V(1/2), 7UDH(1/2), 7UBR(1/2), 7U60(2) | $\alpha_{IIb}\beta_3$ headpiece bound to: sibrafiban, lamifiban, fradafiban, gantofiban, UR-2922, UR-2922, BMS4, BMS4, BMS4.1, BMS4.3, GR144053, cyclic RGDfV, respectively. | Bent-closed state. | $\beta_{Propeller}$, $\beta_A$, Hybrid, PSI. | Lin et al. [S31] |
| 7TD8(1/2), 7THO(1/2), 7UKO(1/2), 7UK9(1/2), 7UFH(1/2), 7UH8(1/2), 7UDG(1/2), 7UKP(1/2), 7TPD(1/2), 7UKT(1/2), 7U60(1) | $\alpha_{IIb}\beta_3$ headpiece bound to: tirofiban, eptifibatide, sibrafiban, lamifiban, fradafiban, roxifiban, lotrafiban, gantofiban analog, EF-5154, BMS4.2, cyclic RGDfV, respectively. | 4 Intermediate states. | $\beta_{Propeller}$, $\beta_A$, Hybrid, PSI. | Lin et al. [S31] |

Table S4: Overview of $\alpha_{IIb}\beta_3$ integrin structures resolved using cryo-EM.

| PDB ID | Description | State | Resolved domains | Reference |
|---|---|---|---|---|
| 6V4P | $\alpha_{IIb}\beta_3$ headpiece bound to eptifibatide. | Bent-closed state. | $\beta_{Propeller}$, $\beta_A$, Hybrid | Nešić et al.[S32] |
| 7LA4 | $\alpha_{IIb}\beta_3$ headpiece bound to PT25-2. | Bent-closed state. | $\beta_{Propeller}$, $\beta_A$, Hybrid | Nešić et al.[S33] |
| 8GCD | Full-length $\alpha_{IIb}\beta_3$. | Bent-closed state. | $\beta_{Propeller}$, Thigh, Calf-1, Calf-2 and $\alpha_{helix}$,$\beta_A$, Hybrid, PSI, I-EGF 1-4, $\beta TD$, and $\beta_{helix}$. | Huo et al.[S34] |
| 8GCE | Ectodomain of $\alpha_{IIb}\beta_3$. | Intermediate state. | $\beta_{Propeller}$, Thigh, Calf-1, Calf-2, $\beta_A$, Hybrid, PSI, I-EGF 1-4, $\beta TD$ | Huo et al.[S34] |
| 8T2U, 8T2V | Full-length $\alpha_{IIb}\beta_3$ bound to: eptifibatide, none, respectively. | Bent-closed state. | $\beta_{Propeller}$, Thigh, Calf-1, Calf-2 and $\alpha_{helix}$, $\beta_A$, Hybrid, PSI, I-EGF 1-4, $\beta TD$, and $\beta helix$. | Adair et al.[S35] |

Table S5: Overview of $\alpha_{IIb}\beta_3$ integrin structures resolved using NMR spectroscopy.

| PDB ID[2] | Description | State | Resolved domains[3] | Reference |
|---|---|---|---|---|
| 1M8O(1-20) | Cytoplasmic domains of $\alpha_{IIb}\beta_3$. | N/A | $\alpha_{helix}$ and $\beta_{helix}$. | Vinogradova et al.[S36] |
| 1DPK(1-16), 1DPQ(1-11) | Wild-type and mutated cytoplasmic domain of $\alpha_{IIb}$, respectively | N/A | $\alpha_{helix}$ | Vinogradova et al.[S37] |
| 1KUP(1-20), 1KUZ(1-20) | Cytoplasmic tail of $\alpha_{IIb}\beta_3$ integrin. | N/A | $\alpha_{helix}$ and $\beta_{helix}$ | Weljie et al.[S38] |
| 1S4W(1-20) | Cytoplasmic domain of $\alpha_{IIb}$. | N/A | $\alpha_{helix}$ | Vinogradova et al.[S39] |
| 2K1A(1-21) | Transmembrane segment of $\alpha_{IIb}$. | N/A. | $\alpha_{helix}$. | Lau et al.[S40] |
| 2K9J(1-21) | $\alpha_{IIb}\beta_3$ transmembrane complex. | N/A. | $\alpha_{helix}$ and $\beta_{helix}$. | Lau et al.[S41] |
| 2KNC(1-20) | $\alpha_{IIb}\beta_3$ transmembrane cytoplasmic domains | N/A | $\alpha_{helix}$ and $\beta_{helix}$. | Yang et al.[S42] |
| 2MTP(1-20) | Filamin-A bound to $\alpha_{IIb}\beta_3$ cytoplasmic domain. | N/A | $\alpha_{helix}$ and $\beta_{helix}$. | Liu et al.[S43] |
| 2N9Y(1-21) | $\alpha_{IIb}\beta_3$ transmembrane complex. | N/A | $\alpha_{helix}$ and $\beta_{helix}$. | Schmidt et al.[S44] |
| 7SFT(1-20) | Filamin-A bound to $\alpha_{IIb}$ cytoplasmic tail. | N/A | $\alpha_{helix}$. | Liu et al.[S45] |
| 7KN0(1-21) | Mutant $\alpha_{IIb}\beta_3$ transmembrane complex with mutation W968V in $\alpha_{IIb}$ subunit. | N/A | $\alpha_{helix}$ and $\beta_{helix}$. | Situ et al.[S46] |

[2] Numbers within braces refers to the model number present in the PDB structure file. For example, (1-20) refers to the indices of the twenty models resolved using NMR spectroscopy.

[3] Only some of the residues within the listed domains are resolved. These are either parts of the residues within the cytoplasmic domains or those in the transmembrane region.

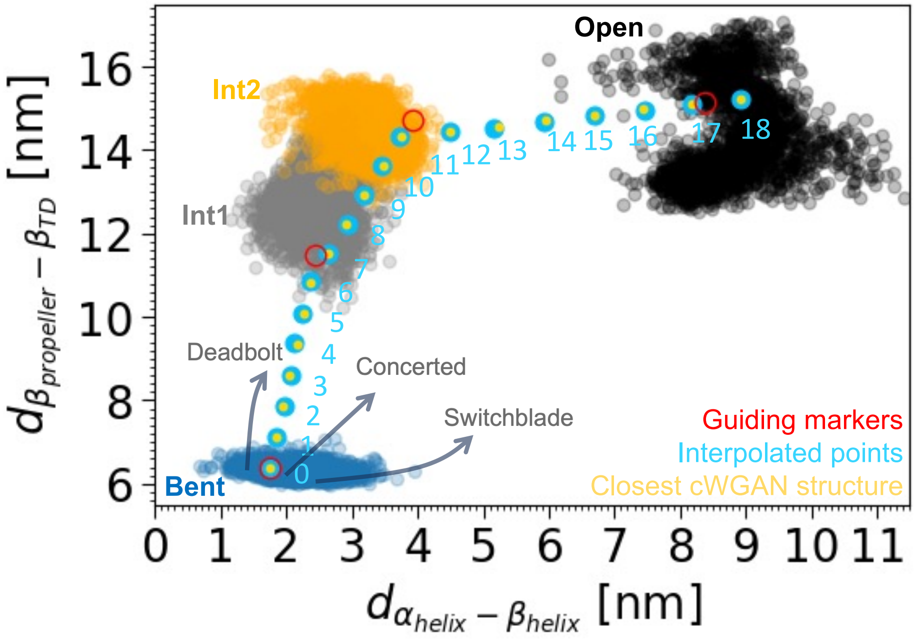


Figure S1: Construction of the initial guess deadbolt activation pathway in the 2D physical CV space spanned by $d_{\alpha_{helix}-\beta_{helix}}$ (x-axis) and $d_{\beta_{propeller}-\beta_{TD}}$ (y-axis). The dark blue, gray, orange, and black markers correspond to the projection of the 2450 MD configurations sampled in Tong et al.[S9] with initial structures resolved using cryo-EM method[S8] in the bent-closed, Int1, Int2, and the extended-open states in the 2D physical CV space, respectively. The equidistant images at 0.75 nm distance along the initial string from bent-closed to extended-open states through the extended intermediates states are shown using light blue markers. These images are constructed by performing three linear interpolations using four manually selected guiding markers shown in red within each state. Specifically, the guiding markers located in bent-closed state at $[1.748, 6.376]$ corresponding to image $\lambda = 0$ and Int1 state at $[2.451, 11.496]$ are first used for defining a unit normal from the bent-closed state to the Int1 state. Then, the equidistant images from $\lambda = 0$ to $\lambda = 6$ are obtained by placing points along this unit normal at 0.75 nm distance. The last image $\lambda = 6$ and the guiding marker at $[3.915, 14.736]$ in the Int2 state are used for constructing the second unit normal from the Int1 state to the Int2 state. This unit normal is then used for determining the images $\lambda = 7$ to $\lambda = 11$ at 0.75 nm distance. Similarly, a third unit normal is constructed using the $\lambda = 11$ and the guiding marker at $[8.374, 15.139]$ in the extended-open state. This unit normal is used for determining the images $\lambda = 12$ to $\lambda = 18$ at 0.75 nm. The locations of the closest available 300 bead CG configuration of $\alpha_{IIb}\beta_3$ integrin from the 90,000 configurations generated by using our multiscale data-driven framework[S1] within this CV space are shown using yellow markers. The arrows illustrate potential traces of the deadbolt, concerted, and switchblade mechanism models in this CV space.

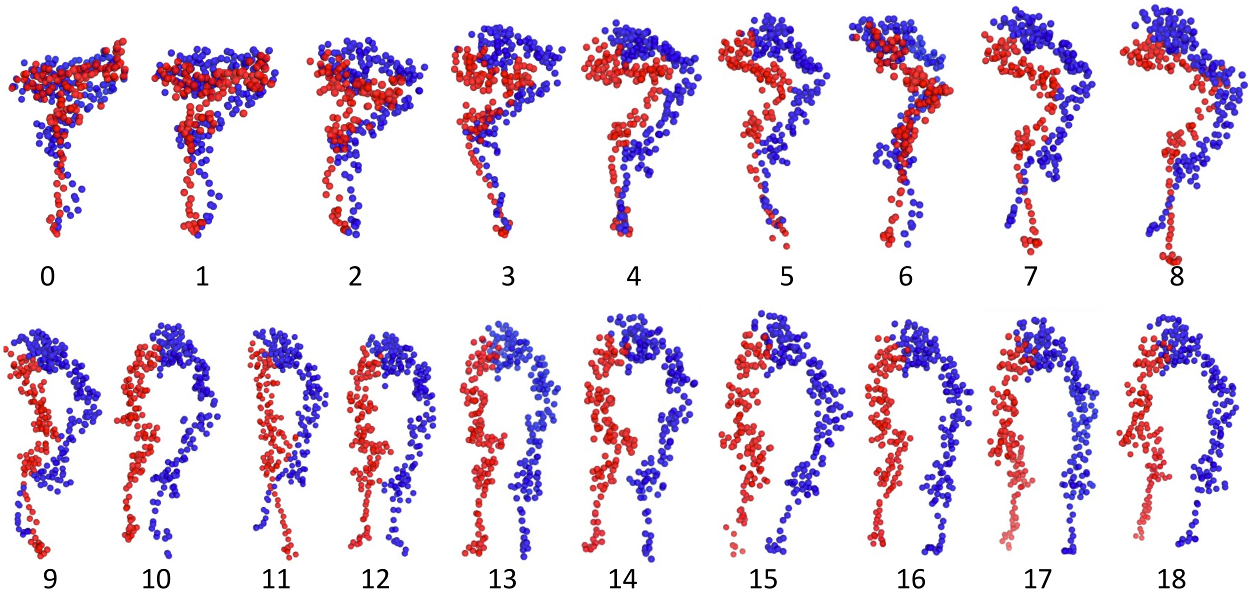


Figure S2: Illustrations of the 300 bead CG configurations corresponding to the yellow markers in SI Fig. S1 that are generated by using our multiscale data-driven framework.[S1] The number below each snapshot corresponds to the image number $\lambda$ along the string. Beads in the $\alpha_{IIb}$ subunit are colored blue, and those in the $\beta_3$ subunit are colored red. The structure along the string demonstrates gradual separation of the $\alpha_{IIb}$ and $\beta_3$ subunits. The structure corresponding to $\lambda = 11$ exhibits a crossing of the tail helices, which, while physically possible, is topologically inconsistent with the neighboring images $\lambda = 10$ and $\lambda = 12$ in the initial path. The physical CVs $d_{\alpha_{helix}-\beta_{helix}}$ (x-axis) and $d_{\beta_{propeller}-\beta_{TD}}$ do not depend strongly on the state of the tail helices, so the path remains smooth on the 2D physical CV space spanned by these two CVs, but this topological inconsistency does introduce an inconsistency into the path trajectory on the CG and AA space that we ameliorate by resetting our pathway through the FTS relaxation process at iteration 249.

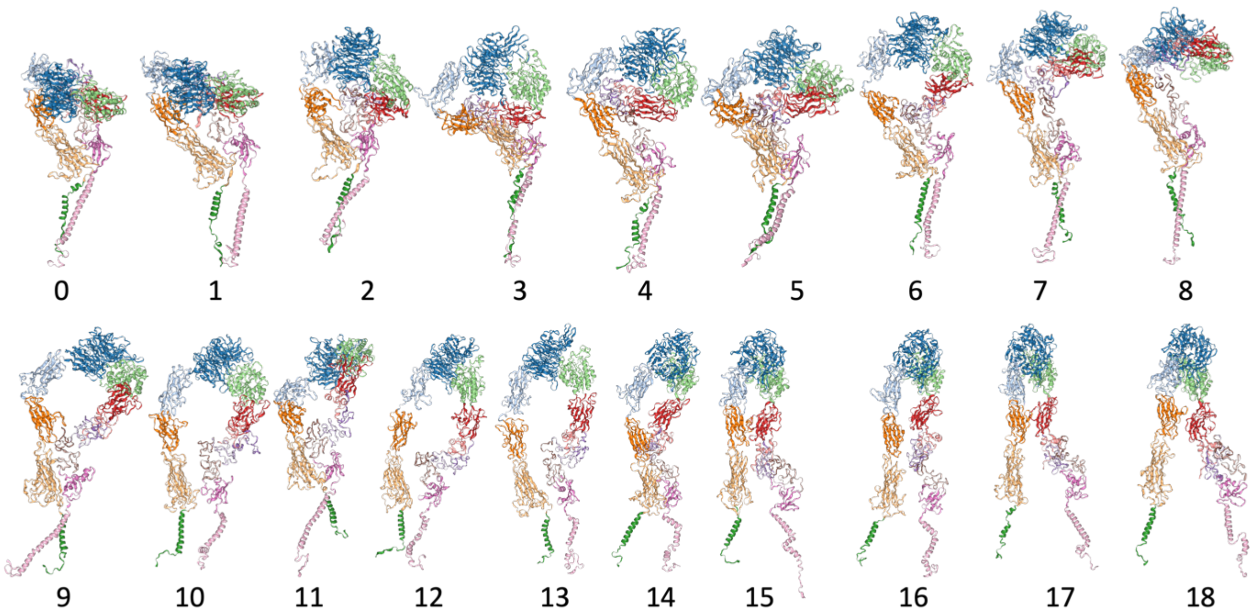


Figure S3: Illustrations of the AA configurations in cartoon representation corresponding to the 300 bead CG configurations in SI Fig. S2. These AA configurations were generated using the TMD simulations (SI Sec.1) and employed to seed the FTS relaxation procedure. The number below each snapshot correspond to the image number $\lambda$ along the string. Each domain in the $\alpha_{IIb}\beta_3$ integrin is colored uniquely. Atoms in lipid bilayer and solvent are not shown in these renderings for clarity. Note the unphysical transition in the tail helices arrangement in the structure corresponding to the $\lambda = 1$ and $\lambda = 11$, where the tail helices fail to follow the transition observed in their neighboring images.

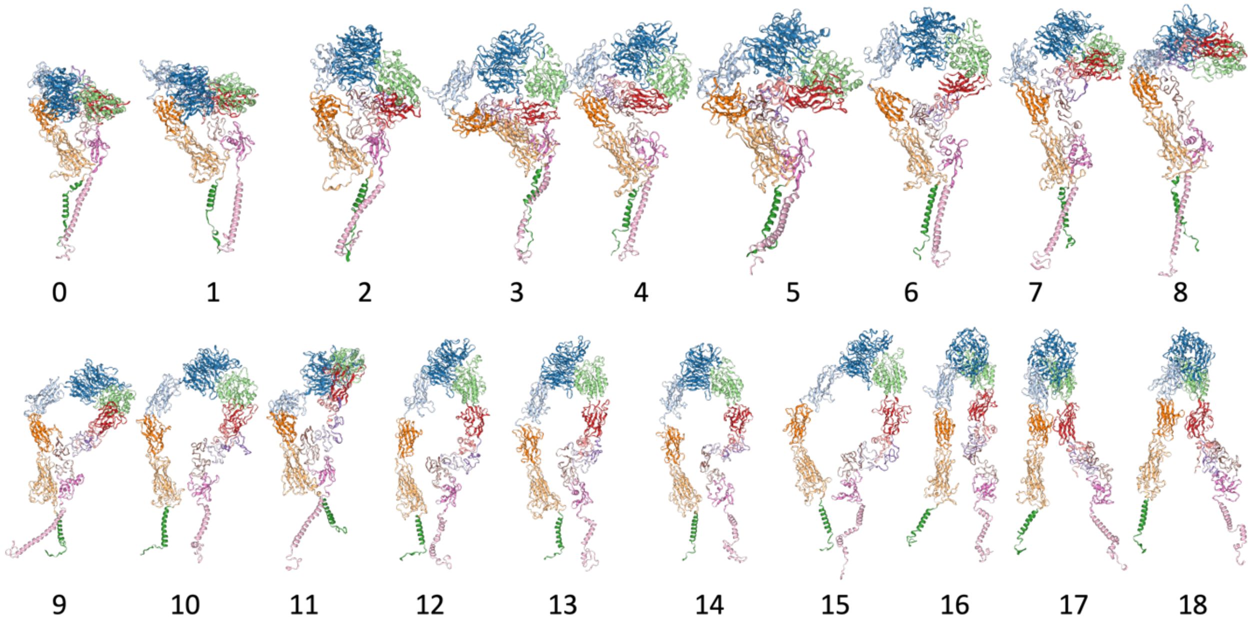


Figure S4: Illustrations of the AA configurations in cartoon representation corresponding to the relaxed string, iteration 249, from the first FTS cycle before re-generation of the structures along the string. The number below each snapshot correspond to the image number $\lambda$ along the string. Each domain in the $\alpha_{IIb}\beta_3$ integrin is colored uniquely. Atoms in lipid bilayer and solvent are not shown in these renderings for clarity.

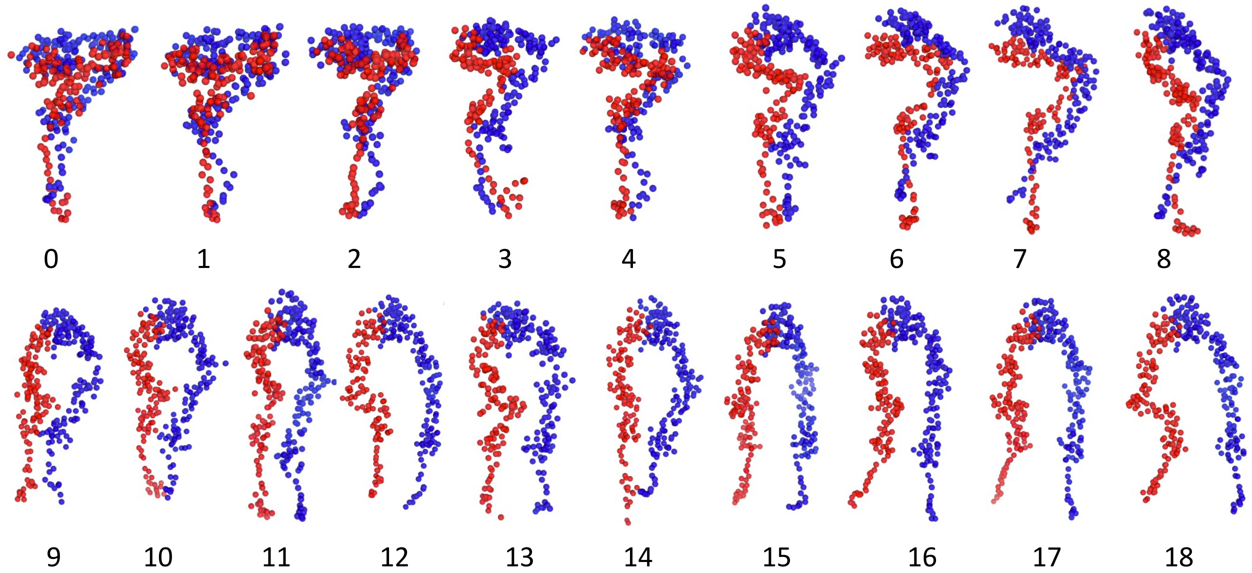


Figure S5: Illustrations of the re-generated 300 bead CG configurations corresponding to the relaxed string, iteration 249, to seed additional round of FTS method cycle. Beads in $\alpha_{IIb}$ subunit and $\beta_3$ subunit are colored in blue and red, respectively. These re-generated images display a smoother transition in all the domains including the tail helices arrangement.

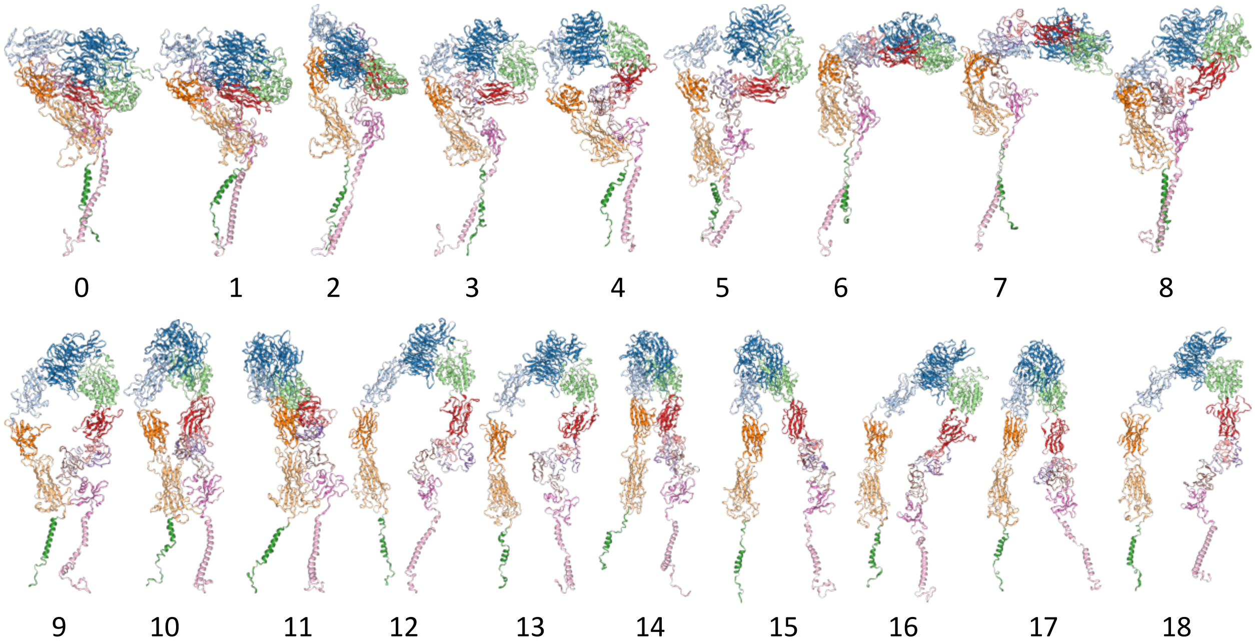


Figure S6: Illustrations of the AA configurations in cartoon representation corresponding to the 300 bead CG configurations in SI Fig. S5. These AA configurations were generated using the TMD simulations and employed to seed the additional round of the FTS method cycle. The number below each snapshot correspond to the image number $\lambda$ along the string. Each domain in the $\alpha_{IIb}\beta_3$ integrin is colored uniquely. Atoms in lipid bilayer and solvent are not shown in these renderings for clarity.

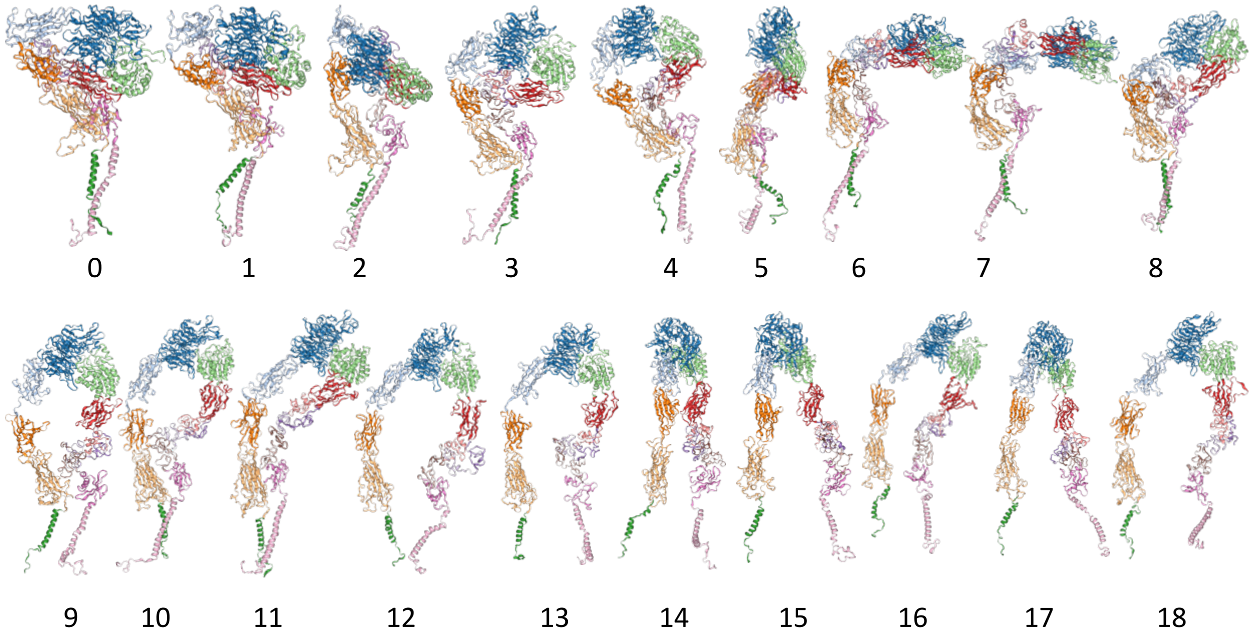


Figure S7: Illustrations of the AA configurations in cartoon representation corresponding to the final relaxed path, iteration 395, from the FTS relaxation process. The number below each snapshot correspond to the image number $\lambda$ along the string. Each domain in the $\alpha_{IIb}\beta_3$ integrin is colored uniquely. Atoms in lipid bilayer and solvent are not shown in these renderings for clarity.

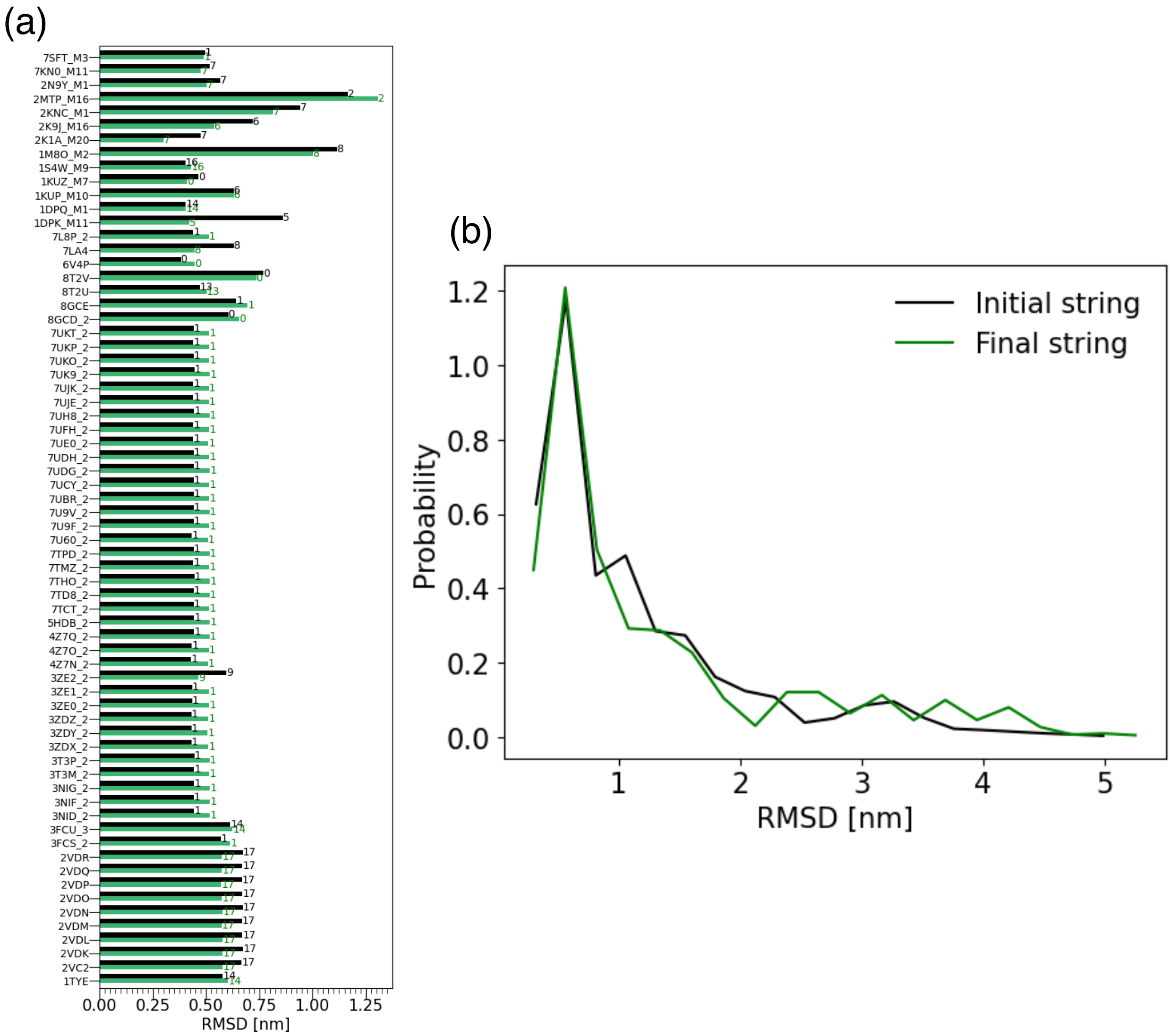


Figure S8: Comparison of the $\alpha_{IIb}\beta_3$ integrin structures along the initial guess pathway (iter 0) and the final (iter 395) relaxed pathway with experimentally resolved structures. (a) RMSD between the structures from the initial (black) relaxed pathway (green) and the structures resolved using X-ray crystallography, NMR spectroscopy, and cryo-EM techniques. The PDB IDs of the experimentally resolved structures are shown along y-axis and their RMSD values with a relaxed structure corresponding to an image, $\lambda$ from the relaxed path that has the least RMSD value are shown along x-axis. The image number $\lambda$ with the minimum RMSD from the experimentally resolved structures are shown on the bar. (b) Distribution of the RMSD between all the structures along the initial (black) and the final (green) relaxed pathway with all the structures resolved using experimental methods, including the multiple structures resolved in NMR.

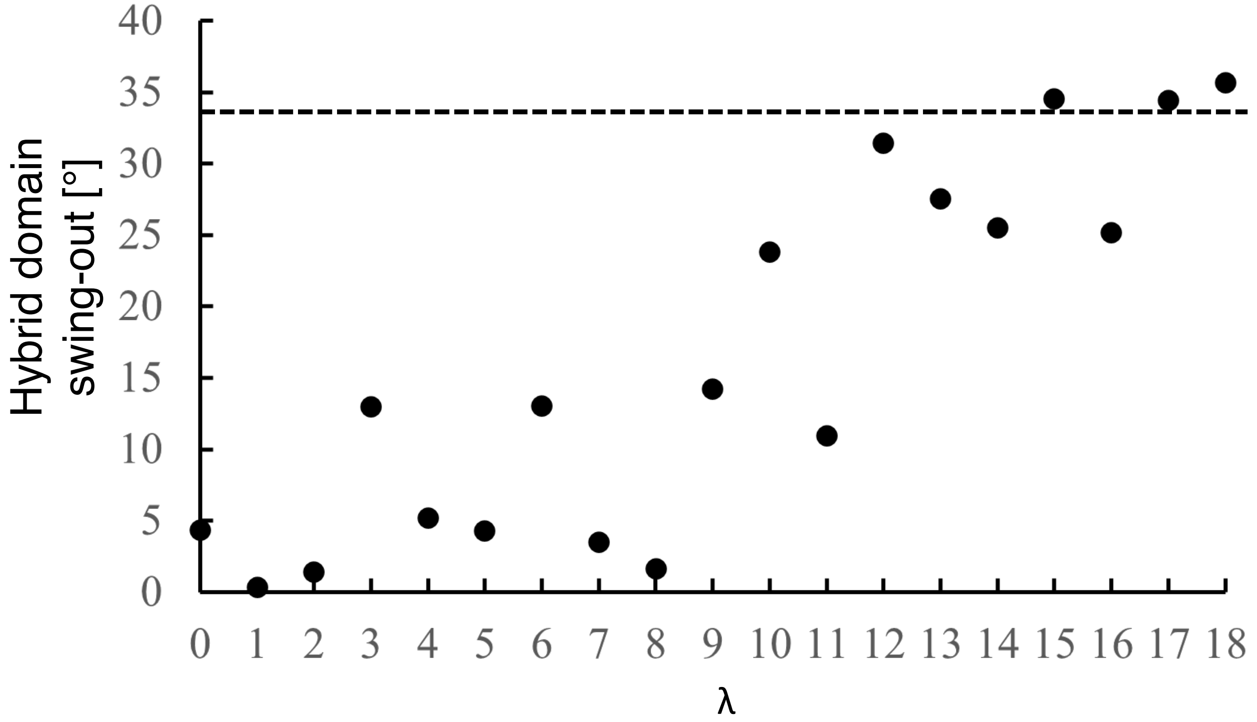


Figure S9: Changes in the hybrid domain swing-out angle along the final relaxed string (iter 395) from the FTS relaxation process. We define the hybrid domain's swing-out angle as the angle between the vector formed by the COM of the hybrid domain and the $\beta_A$ domain in the structure corresponding to an image and the same vector in a reference experimentally (X-ray crytallography) resolved structure with PDB: 3T3P. The changes in hybrid domain's swing-out angle remains relatively level and low from $\lambda = 1$ through $\lambda = 8$. It follows with images $\lambda = 9 - 11$ that appear to be in a transition state, and the images $\lambda = 12 - 18$ transition into a completely swung-out or the extended-open state. All the structures corresponding to the images along the final relaxed string (iter 395) from the FTS relaxation process are first aligned to the reference structure using $C_\alpha$ atoms in the $\beta_A$ domain prior to the estimation of the hybrid domain's swing-out angle. For reference on the hybrid-domain swung-out angle, we show with horizontal dotted line the angle formed by the vector in the experimental structures PDB: 3T3P and PDB: 2VDR that correspond to the bent-closed state and the extended-open state, respectively.

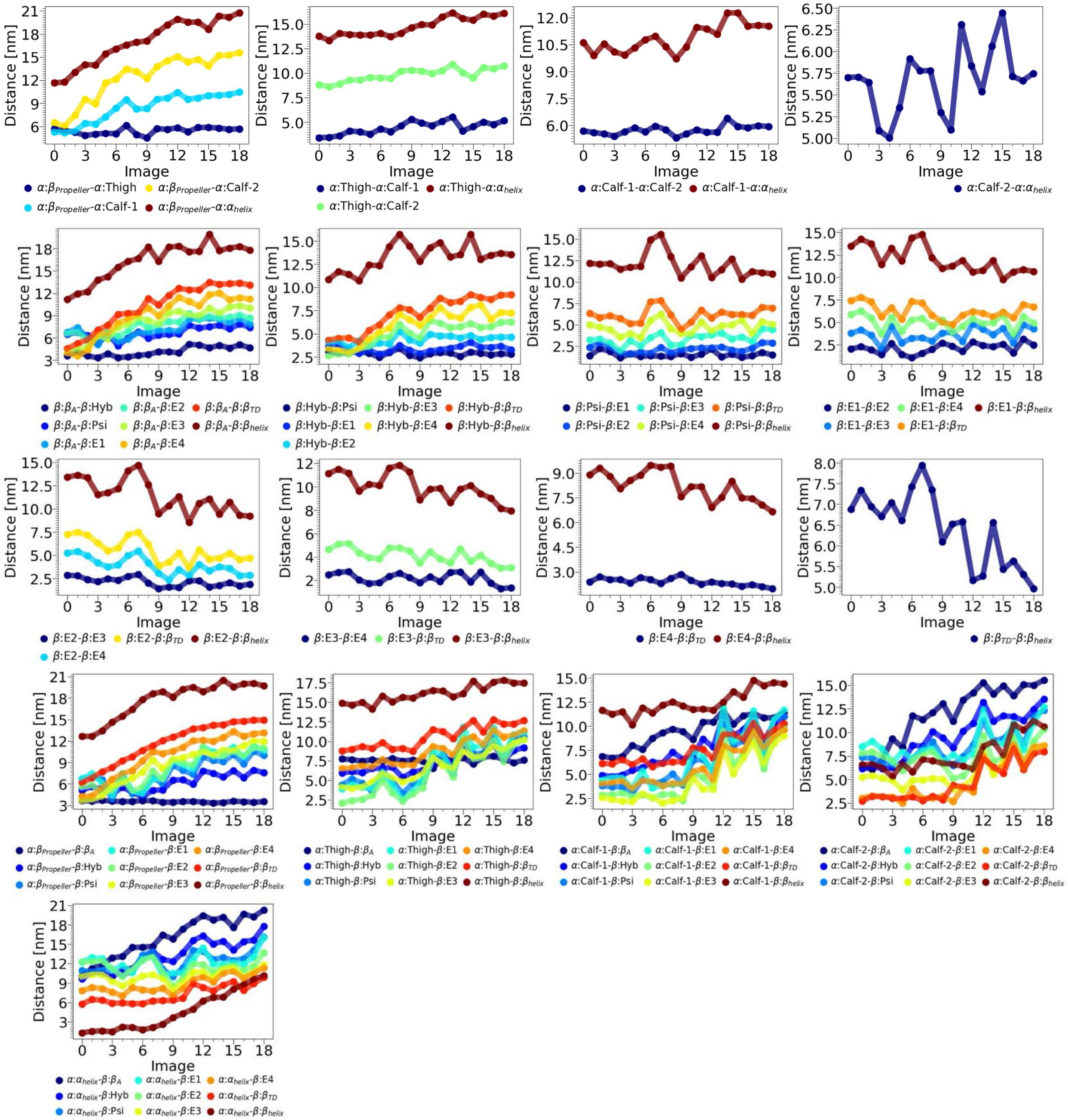

Figure S10: Changes in the COM distance between different domains in the structures along the final relaxed path, iteration 395, from the FTS relaxation process.

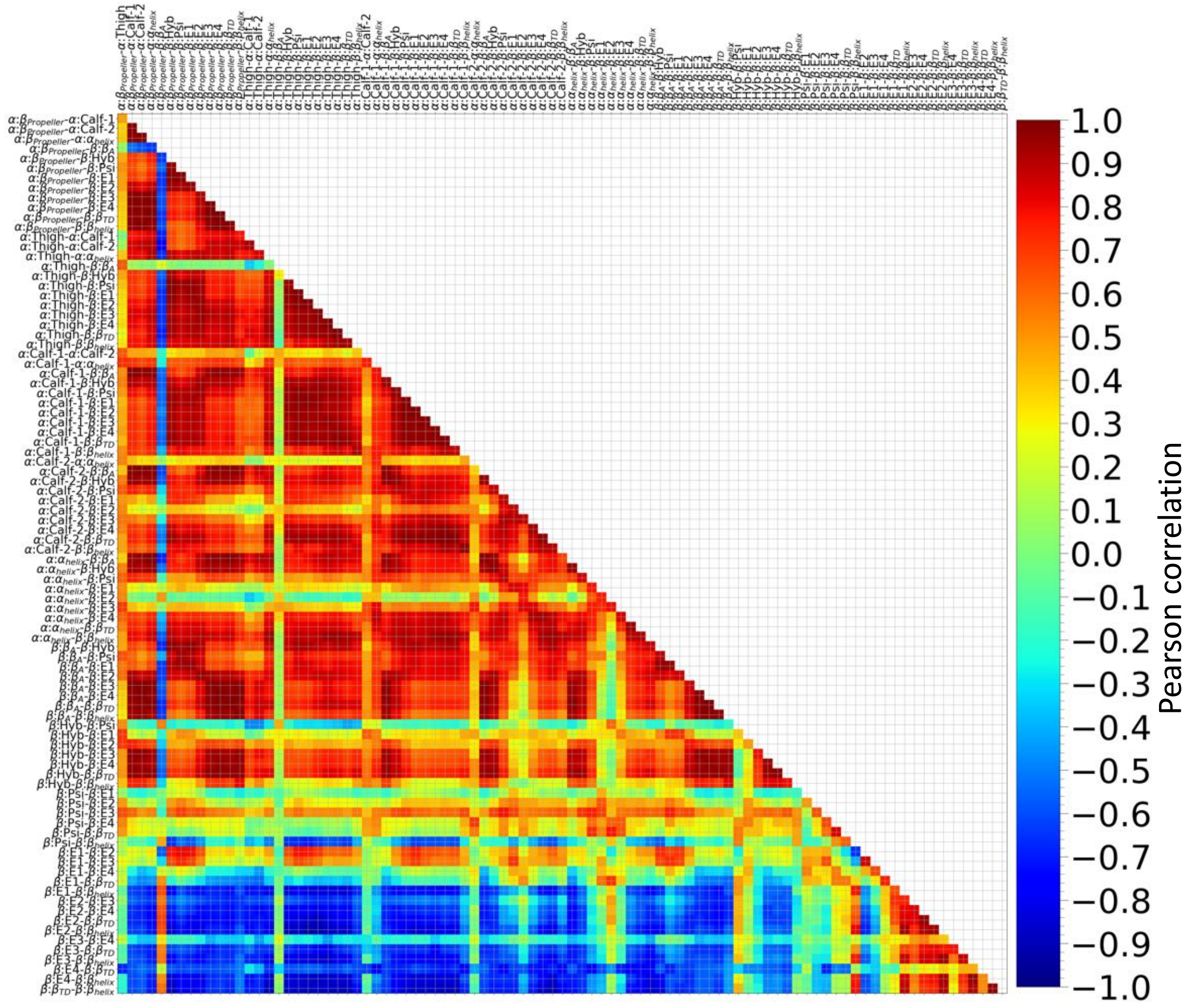


Figure S11: Pearson correlation between the change in the COM distances shown in SI Fig. S10 between different domains in the structures along the final relaxed path, iteration 395, from the FTS relaxation process. For example, the top left pixel with a light orange shade shows the Pearson correlation between the changes in the COM distance between the domains $\beta_{propeller}$ and Calf-1 in $\alpha$ subunit ($\alpha$ : $\beta_{propeller} - \alpha$ :Calf-1) and the changes in the COM distance between the domains $\beta_{propeller}$ and Thigh domain in $\alpha$ subunit ($\alpha$ : $\beta_{propeller} - \alpha$ :Thigh) along the relaxed path, iteration 395.

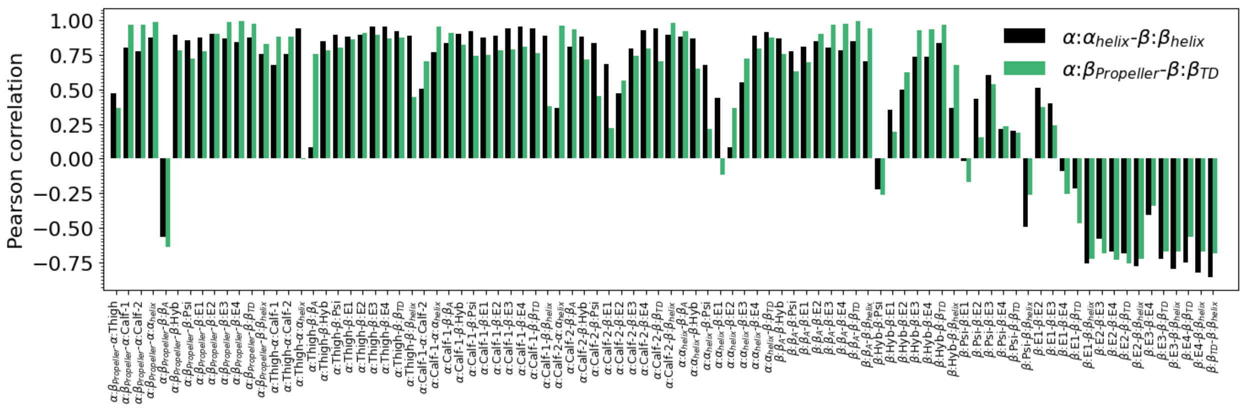


Figure S12: Pearson correlations between the changes in the CVs ($d_{\alpha_{helix}-\beta_{helix}}$, $d_{\beta_{propeller}-\beta_{TD}}$) with change in the COM distances between other domain pairs in $\alpha_{IIb}\beta_3$ integrin along the relaxed path (iter 395) from the second relaxation process. The Pearson correlations with $d_{\alpha_{helix}-\beta_{helix}}$ and $d_{\beta_{propeller}-\beta_{TD}}$) are shown using black and green color bars, respectively. These Pearson correlations correspond to the columns $d_{\alpha_{helix}-\beta_{helix}}$ and $d_{\beta_{propeller}-\beta_{TD}}$ in the symmetric matrix plot shown in SI Fig. S11

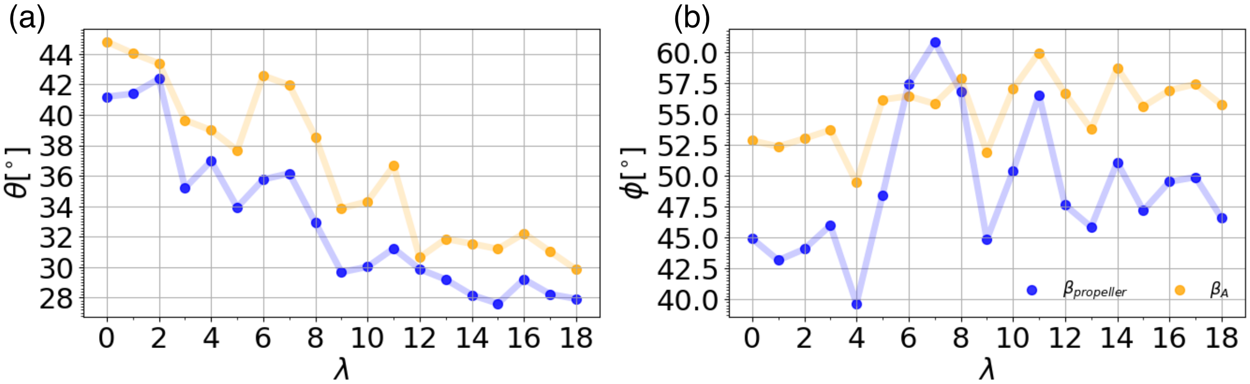


Figure S13: Changes in the orientation of the headpiece of $\alpha_{IIb}\beta_3$ integrin along the final relaxed string (iter 395) from the FTS relaxation process. Variations of (a) polar ($\theta$) and (b) azimuthal ($\phi$) angles of $\beta_{propeller}$ and $\beta_A$ domains of $\alpha_{IIb}\beta_3$ integrin along the relaxed string from the FTS method (Fig. 5 in main text). The structures corresponding to $\lambda = 1, ..18$ are aligned in the plane of the lipid bilayer to the structure of the image $\lambda = 0$ before converting the Cartesian coordinates of the COM of each domain to the spherical coordinate system.

(S5) Gulrajani, I.; Ahmed, F.; Arjovsky, M.; Dumoulin, V.; Courville, A. C. Improved Training of Wasserstein GANs. Advances in Neural Information Processing Systems. 2017.

(S6) Schlitter, J.; Engels, M.; Krüger, P. *J. Mol. Graphics* **1994**, *12*, 84–89.

(S7) Hénin, J.; Lelièvre, T.; Shirts, M.; Valsson, O.; Delemotte, L. *Living J. Comput. Mol. Sci.* **2022**, *4*, 1583–1583.

(S8) Xu, X.-P.; Kim, E.; Swift, M.; Smith, J. W.; Volkmann, N.; Hanein, D. *Biophys. J.* **2016**, *110*, 798–809.

(S9) Tong, D.; Soley, N.; Kolasangiani, R.; Schwartz, M. A.; Bidone, T. C. *Biophys. J.* **2023**, *122*, 533–543.

(S10) Mongan, J.; Simmerling, C.; McCammon, J. A.; Case, D. A.; Onufriev, A. *J. Chem. Theory Comput.* **2007**, *3*, 156–169.

(S11) Best, R. B.; Zhu, X.; Shim, J.; Lopes, P. E.; Mittal, J.; Feig, M.; MacKerell Jr, A. D. *J. Chem. Theory Comput.* **2012**, *8*, 3257–3273.

(S12) Eastman, P.; Pande, V. S. *J. Chem. Theory Comput.* **2010**, *6*, 434–437.

(S13) Eastman, P.; Friedrichs, M. S.; Chodera, J. D.; Radmer, R. J.; Bruns, C. M.; Ku, J. P.; Beauchamp, K. A.; Lane, T. J.; Wang, L.-P.; Shukla, D.; Tye, T.; Houston, M.;

Stich, T.; Klein, C.; Shirts, M. R.; Pande, V. S. *J. Chem. Theory Comput.* **2013**, *9*, 461–469.

(S14) Tribello, G. A.; Bonomi, M.; Branduardi, D.; Camilloni, C.; Bussi, G. *Comput. Phys. Commun.* **2014**, *185*, 604–613.

(S15) Jo, S.; Kim, T.; Iyer, V. G.; Im, W. *J. Comput. Chem.* **2008**, *29*, 1859–1865.

(S16) Jo, S.; Kim, T.; Im, W. *PloS One* **2007**, *2*, e880.

(S17) Lee, J.; Cheng, X.; Swails, J. M.; Yeom, M. S.; Eastman, P. K.; Lemkul, J. A.; Wei, S.; Buckner, J.; Jeong, J. C.; Qi, Y.; Jo, S.; Pande, V. S.; Case, D. A.; Brooks, C. L. I.; MacKerell, A. D. J.; Klauda, J. B.; Im, W. *Journal of Chemical Theory and Computation* **2016**, *12*, 405–413, PMID: 26631602.

(S18) Berendsen, H. J. C.; Postma, J. P. M.; van Gunsteren, W. F.; DiNola, A.; Haak, J. R. *J. Chem. Phys.* **1984**, *81*, 3684–3690.

(S19) Abraham, M. J.; Murtola, T.; Schulz, R.; Páll, S.; Smith, J. C.; Hess, B.; Lindahl, E. *SoftwareX* **2015**, *1*, 19–25.

(S20) Lindahl, E.; Abraham, M.; Hess, B.; van der Spoel, D. GROMACS 2019.3 Source code. `https://doi.org/10.5281/zenodo.3243833`, 2019; `https://doi.org/10.5281/zenodo.3243833`.

(S21) Darden, T.; York, D.; Pedersen, L. *J. Chem. Phys.* **1993**, *98*, 10089–10092.

(S22) Hess, B.; Bekker, H.; Berendsen, H. J. C.; Fraaije, J. G. E. M. *J. Comput. Chem.* **1997**, *18*, 1463–1472.

(S23) Verlet, L. *Phys. Rev.* **1967**, *159*, 98–103.

(S24) Xiao, T.; Takagi, J.; Coller, B. S.; Wang, J.-H.; Springer, T. A. *Nature* **2004**, *432*, 59–67.

(S25) Springer, T. A.; Zhu, J.; Xiao, T. *J. Cell Biol.* **2008**, *182*, 791–800.

(S26) Zhu, J.; Luo, B.-H.; Xiao, T.; Zhang, C.; Nishida, N.; Springer, T. A. *Mol. Cell* **2008**, *32*, 849–861.

(S27) Zhu, J.; Zhu, J.; Negri, A.; Provasi, D.; Filizola, M.; Coller, B. S.; Springer, T. A. *Blood* **2010**, *116*, 5050–5059.

(S28) Zhu, J.; Choi, W.-S.; McCoy, J. G.; Negri, A.; Zhu, J.; Naini, S.; Li, J.; Shen, M.; Huang, W.; Bougie, D.; Rasmussen, M.; Aster, R.; Thomas, C. J.; Filizola, M.; Springer, T. A.; Coller, B. S. *Sci. Transl. Med.* **2012**, *4*, 125ra32–125ra32.

(S29) Zhu, J.; Zhu, J.; Springer, T. A. *J. Cell Biol.* **2013**, *201*, 1053–1068.

(S30) Lin, F.-Y.; Zhu, J.; Eng, E. T.; Hudson, N. E.; Springer, T. A. *Journal of Biol. Chem.* **2016**, *291*, 4537–4546.

(S31) Lin, F.-Y.; Li, J.; Xie, Y.; Zhu, J.; Nguyen, T. T. H.; Zhang, Y.; Zhu, J.; Springer, T. A. *Cell* **2022**, *185*, 3533–3550.

(S32) Nešić, D.; Zhang, Y.; Spasic, A.; Li, J.; Provasi, D.; Filizola, M.; Walz, T.; Coller, B. S. *Arterioscler. Thromb. Vasc. Biol.* **2020**, *40*, 624–637.

(S33) Nešić, D.; Bush, M.; Spasic, A.; Li, J.; Kamata, T.; Handa, M.; Filizola, M.; Walz, T.; Coller, B. S. *Blood Adv.* **2021**, *5*, 1781–1790.

(S34) Huo, T.; Wu, H.; Moussa, Z.; Sen, M.; Dalton, V.; Wang, Z. *Structure* **2024**,

(S35) Adair, B. D.; Xiong, J.-P.; Yeager, M.; Arnaout, M. A. *Nat. Comm.* **2023**, *14*, 4168.

(S36) Vinogradova, O.; Velyvis, A.; Velyviene, A.; Hu, B.; Haas, T. A.; Plow, E. F.; Qin, J. *Cell* **2002**, *110*, 587–597.

(S37) Vinogradova, O.; Haas, T.; Plow, E. F.; Qin, J. *Proc. Natl. Acad. Sci. U.S.A.* **2000**, *97*, 1450–1455.

(S38) Weljie, A. M.; Hwang, P. M.; Vogel, H. J. *Proc. Natl. Acad. Sci. U.S.A.* **2002**, *99*, 5878–5883.

(S39) Vinogradova, O.; Vaynberg, J.; Kong, X.; Haas, T. A.; Plow, E. F.; Qin, J. *Proc. Natl. Acad. Sci. U.S.A.* **2004**, *101*, 4094–4099.

(S40) Lau, T.-L.; Dua, V.; Ulmer, T. S. *J. Biol. Chem.* **2008**, *283*, 16162–16168.

(S41) Lau, T.-L.; Kim, C.; Ginsberg, M. H.; Ulmer, T. S. *EMBO J.* **2009**, *28*, 1351–1361.

(S42) Yang, J.; Ma, Y.-Q.; Page, R. C.; Misra, S.; Plow, E. F.; Qin, J. *Proc. Natl. Acad. Sci. U.S.A.* **2009**, *106*, 17729–17734.

(S43) Liu, J.; Das, M.; Yang, J.; Ithychanda, S. S.; Yakubenko, V. P.; Plow, E. F.; Qin, J. *Nat. Struct. Mol. Biol.* **2015**, *22*, 383–389.

(S44) Schmidt, T.; Situ, A. J.; Ulmer, T. S. *Sci. Rep.* **2016**, *6*, 29809.

(S45) Liu, J.; Lu, F.; Ithychanda, S. S.; Apostol, M.; Das, M.; Deshpande, G.; Plow, E. F.; Qin, J. *Blood* **2023**, *141*, 2629–2641.

(S46) Situ, A. J.; Kim, J.; An, W.; Kim, C.; Ulmer, T. S. *J. Mol. Biol.* **2021**, *433*, 166832.
